# Human gene function publications that describe wrongly identified nucleotide sequence reagents are unacceptably frequent within the genetics literature

**DOI:** 10.1101/2021.07.29.453321

**Authors:** Yasunori Park, Rachael A West, Pranujan Pathmendra, Bertrand Favier, Thomas Stoeger, Amanda Capes-Davis, Guillaume Cabanac, Cyril Labbé, Jennifer A Byrne

## Abstract

Nucleotide sequence reagents underpin a range of molecular genetics techniques that have been applied across hundreds of thousands of research publications. We have previously reported wrongly identified nucleotide sequence reagents in human gene function publications and described a semi-automated screening tool Seek & Blastn to fact-check the targeting or non-targeting status of nucleotide sequence reagents. We applied Seek & Blastn to screen 11,799 publications across 5 literature corpora, which included all original publications in *Gene* from 2007-2018 and all original open-access publications in *Oncology Reports* from 2014-2018. After manually checking the Seek & Blastn screening outputs for over 3,400 human research papers, we identified 712 papers across 78 journals that described at least one wrongly identified nucleotide sequence. Verifying the claimed identities of over 13,700 nucleotide sequences highlighted 1,535 wrongly identified sequences, most of which were claimed targeting reagents for the analysis of 365 human protein-coding genes and 120 non-coding RNAs, respectively. The 712 problematic papers have received over 17,000 citations, which include citations by human clinical trials. Given our estimate that approximately one quarter of problematic papers are likely to misinform or distract the future development of therapies against human disease, urgent measures are required to address the problem of unreliable gene function papers within the literature.

**Author summary:** This is the first study to have screened the gene function literature for nucleotide sequence errors at the scale that we describe. The unacceptably high rates of human gene function papers with incorrect nucleotide sequences that we have discovered represent a major challenge to the research fields that aim to translate genomics investments to patients, and that commonly rely upon reliable descriptions of gene function. Indeed, wrongly identified nucleotide sequence reagents represent a double concern, as both the incorrect reagents themselves and their associated results can mislead future research, both in terms of the research directions that are chosen and the experiments that are undertaken. We hope that our research will inspire researchers and journals to seek out other problematic human gene function papers, as we are unfortunately concerned that our results represent the tip of a much larger problem within the literature. We hope that our research will encourage more rigorous reporting and peer review of gene function results, and we propose a series of responses for the research and publishing communities.

## Introduction

The promise of genomics to improve the health of cancer and other patients has resulted in billions of dollars of research investment which have been accompanied by expectations of similar quantum gains in health outcomes (1, 2). Since the first draft of the human genome was reported (3, 4), a series of increasingly rapid technological advances has permitted the routine sequencing of human genomes at scale (1, 2), and the increasing application of genomics to inform clinical care (1, 2, 5). Despite the now routine capacity to sequence the human genome, genomics research relies upon results produced by other research fields to translate genome sequencing results to patients (5–7). For example, while whole genome sequencing demonstrates that thousands of human genes are mutated or deregulated in human cancers (1), knowledge of human gene function is required to prioritise individual gene candidates for subsequent pre-clinical and translational studies (5–7).

A first step in triaging and prioritising gene candidates for further analysis is the consideration of available knowledge of predicted and/or demonstrated gene functions (5–8). High quality, reliable information about gene function is important to select the most promising gene candidates and to then progress these candidates through pre-clinical and translational research pipelines (8), which is supported by drug candidates with genetically supported targets being significantly more likely to progress through phased clinical trials (9, 10). However, in contrast to the sophisticated platforms that produce genomic or transcriptomic sequence data at scale, gene function experiments typically analyse single or small numbers of genes through the application of more ubiquitous molecular techniques (6), some of which have been in routine experimental use for 15-30 years. For example, gene knockdown approaches have been widely employed to assess the consequences of reduced gene expression in model systems (6). Similarly, RT-PCR is frequently used to analyse the transcript levels of small groups of genes, either to confirm the effectiveness of gene knockdown experiments or in association with other experimental techniques. The widespread use and reporting of gene knockdown and PCR approaches reflect their low cost and accessibility, in terms of the necessary reagents, laboratory equipment and facilities, and the availability of technical expertise within the research community. As a consequence, the results of experiments employing gene knockdown and/or PCR in the context of human research have been described in hundreds of thousands of publications that are retrievable through PubMed or Google Scholar.

Experiments that analyse the functions of individual genes typically require nucleotide sequence reagents as either targeting and/or control reagents, with the sequences corresponding to these reagents being disclosed to indicate the exact experiments that were conducted (8, 11). As nucleotide sequence identities cannot be deduced by eye, DNA or RNA reagent sequences must be paired with text descriptions of their genetic identities and experimental use (8, 11–13). The integrity of reported experiments therefore requires both the identities of nucleotide sequence reagents and their text descriptions to be correct (11, 12). Accurate reporting of nucleotide sequence reagents is also critical to permit reagent reuse across different experiments and publications (11).

The ubiquitous description of nucleotide sequence reagents within the biomedical and genetics literature, combined with the routine pairing of nucleotide sequences and text identifiers, are likely to contribute to tacit assumptions that reported nucleotide sequence reagents are correctly identified. However, as nucleotide sequences cannot be understood by eye, we have proposed that nucleotide sequence reagents are susceptible to different types of errors (8, 11, 12). These error types represent the equivalent of spelling errors (12, 14, 15), as well as identity errors, where a correct sequence is replaced by a different and possibly genetically unrelated sequence (11–13, 16–21).

The problem of wrongly identified nucleotide sequences was recognised in the context of DNA microarrays in the early 2000’s, where wrongly identified sequence probes affected the reliability and reproducibility of data from particular microarray platforms (22, 23). Our team subsequently identified frequent wrongly identified sh/siRNA’s and RT-PCR primers in studies that reported the effects of knocking down single human genes in cancer cell lines (12, 13). Some incorrect sh/siRNA’s invalidated the experimental results reported, through for example the repeated use of “non-targeting” shRNA’s that were in fact indicated to target specific human genes (11–13). Our discovery of incorrect and indeed impossible results associated with wrongly identified nucleotide sequence reagents led us to develop a semi-automated tool Seek & Blastn (S&B) to fact-check the reported identities of nucleotide sequence reagents in human research publications (12, 24). The S&B tool scans text to identify and extract nucleotide sequences and their associated text descriptors, submits extracted nucleotide sequences to blastn analysis (25) to predict their genetic identities and hence their targeting or non-targeting status, and then compares the predicted status of each nucleotide sequence reagent with the claimed status within the text (12). The blastn results for each extracted nucleotide sequence are then reported and any sequence whose text identifier contradicts its blastn-predicted targeting/non-targeting status is flagged as being potentially incorrect (12). Flagged nucleotide sequences are then subjected to manual verification, as described in our original publication (12) and an expanded online protocol (26).

Our initial application of S&B identified 77/203 (38%) screened papers with incorrect nucleotide sequence reagents, with our focus being the description of the S&B tool (12), as opposed to its application. We have now employed S&B to screen original research papers across 5 literature corpora, representing 3 targeted and two journal corpora. The 3 targeted corpora included papers that were identified through literature searches that employed specific keywords, and in some cases, PubMed similarity searches of index papers. The two journal corpora included all original and original open-access papers in *Gene* and *Oncology Reports,* respectively, as examples of journals that have published papers with wrongly identified nucleotide sequences (11–13). S&B screening of 11,799 papers flagged 3,423 papers for further analysis, which required manually verifying the identities of over 13,700 nucleotide sequences. Our combined results provide very worrying evidence that human research papers with wrongly identified nucleotide sequences are unacceptably frequent within the literature.

## Results

### Analysis of targeted publication corpora

To extend our previous results from employing S&B to screen gene function papers (11, 12), we used S&B to screen 3 targeted publication corpora (Fig 1) that were identified through literature searches that employed specific keywords, and in some cases, PubMed similarity searches of index papers (12, 13). In all cases, the keywords that were used to derive targeted corpora did not refer to author affiliations, such as institution type or country of origin (see Methods).

**Fig 1.**
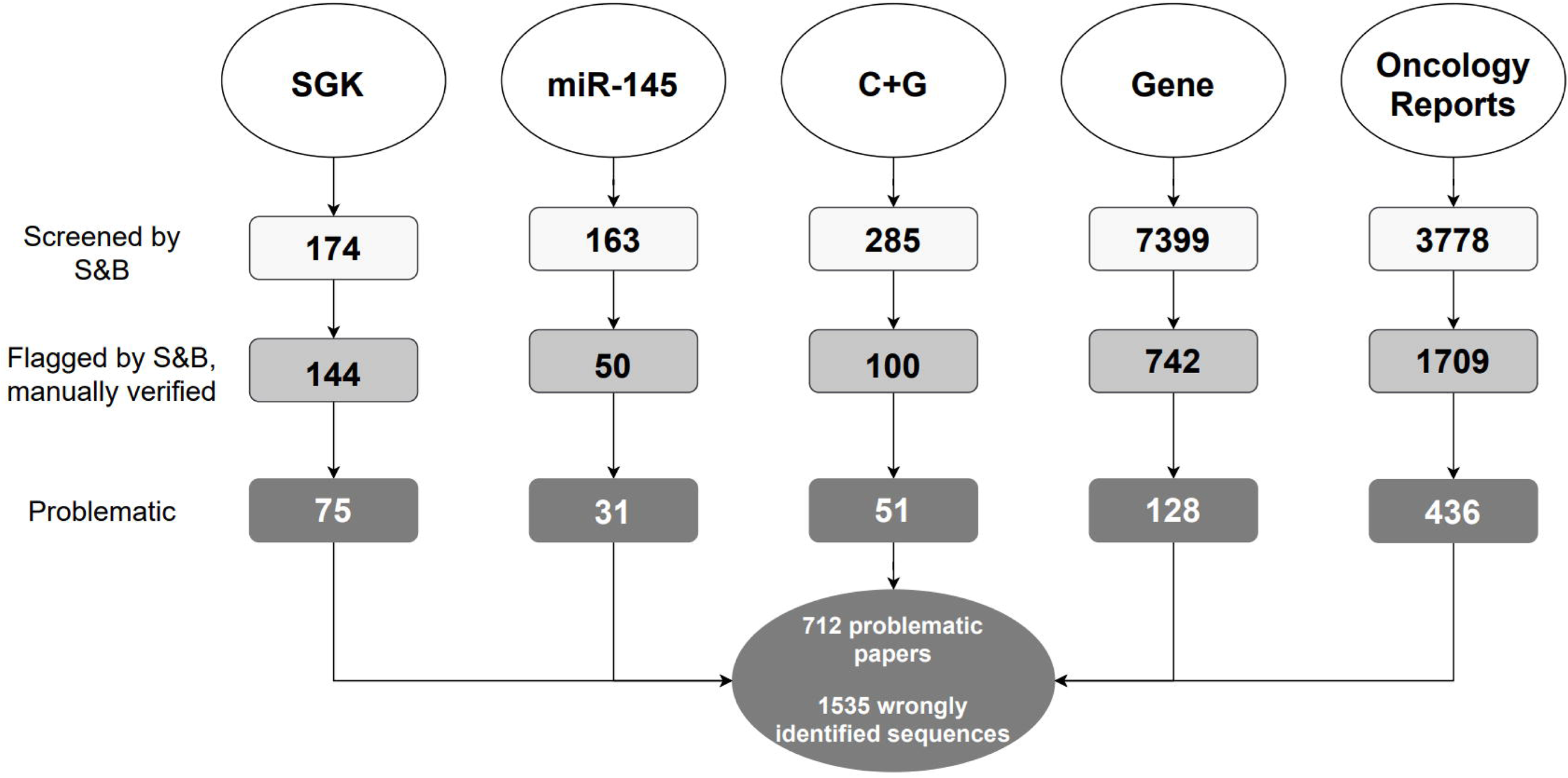
Diagram describing the five literature corpora screened by S&B. For each corpus (top row), the diagram shows the numbers of articles that were (i) screened by S&B (white), (ii) flagged by S&B with sequences manually verified (grey), and (iii) found to be problematic by describing at least one wrongly identified nucleotide sequence (dark grey). Total numbers of problematic papers and wrongly identified sequences are indicated below the diagram, corrected for duplicate papers between the corpora.

#### Single gene knockdown (SGK) corpus

We have previously identified frequent wrongly identified nucleotide sequences in papers that reported the effects of knocking down a single human gene in cancer cell lines that corresponded to a single cancer type, which we referred to as single gene knockdown (SGK) papers (12, 13). As a subset of SGK papers analysed the same human genes across multiple papers and cancer types (12, 13), we sought to identify all SGK papers for a set of 17 human genes (*ADAM8, ANXA1, EAG1, GPR137, ICT1, KLF8, MACC1, MYO6, NOB1, PP4R1, PP5, PPM1D, RPS15A, TCTN1, TPD52L2, USP39,* and *ZFX)*, most of which had been analysed in previously reported papers (12, 13).

By combining each of the 17 human gene identifiers with keywords previously used to identify SGK papers (13), we identified 174 SGK papers published between 2006-2019 across 83 journals (Table 1). As most gene identifiers were selected from previously reported SGK papers (12, 13), the SGK corpus consisted of 41 (24%) previously reported papers and 132 (76%) additional papers (Table 2). All 174 SGK papers analysed gene function in a single human cancer type (Table 2). Across the 17 queried genes, we identified a median of 8 papers/gene (range 3-20) that analysed a median of 8 human cancer types (range 3-11) (Table 2, S1 Data). Most (136/174, 78%) SGK papers named a single queried gene in their titles, with the remaining titles also referring to other human gene(s) and/or drugs, most frequently cisplatin (S1 Data). Most (159/174, 91%) SGK papers were published by authors from mainland China, almost all of which were affiliated with hospitals (147/159, 92%) (see Methods) (Table 1). In contrast, less than half (6/15, 40%) SGK papers from 5 other countries were affiliated with hospitals (Table 1, S1 Data).

**Table 1:** Descriptions of the targeted corpora screened by Seek & Blastn with manual verification of nucleotide sequence reagent identities

|  | Single gene knockdown (SGK) |  | *miR-145* |  | Cisplatin + Gemcitabine (C+G) |  |
| --- | --- | --- | --- | --- | --- | --- |
|  | Corpus | Problematic | Corpus | Problematic | Corpus | Problematic |
| Number of papers<br>(% of corpus) | 174<br>(100%) | **75**<br>**(43%)** | 50<br>(100%) | **31**<br>**(62%)** | 100<br>(100%) | **51**<br>**(50%)** |
| Number of journals | 83 | **42** | 35 | **25** | 48 | **31** |
| Publication year<br>median (range) | 2015<br>(2006-2019)<sup>a</sup> | **2015**<br>**(2010-2019)** | 2017<br>(2009-2019) | **2017**<br>**(2009-2019)** | 2017<br>(2008-2019) | **2017**<br>**(2009-2019)** |
| Journal Impact Factor at publication year<br>median (range) | 2.204<br>(0.098-8.459) | **1.778**<br>**(0.098-5.712)** | 3.34<br>(0.700-9.050) | **3.23**<br>**(0.700-8.278)** | 3.571<br>(1.099-10.391) | **3.041**<br>**(1.099-8.579)** |
| Number of sequences/ paper<br>median (range) | 6 (0-24) | **6 (1-24)** | 11 (4-46) | **10 (4-46)** | 11 (2-71) | **12 (2-70)** |
| Number of incorrect sequences/ paper<br>median (range) | ND | **1 (1-8)** | ND | **1 (1-5)** | ND | **2 (1-8)** |
| Papers from China<br>proportion (%) | 159/174<br>(91%) | **73/75**<br>**(97%)** | 44/50<br>(88%) | **31/31**<br>**(100%)** | 90/100<br>(90%) | **50/51**<br>**(98%)** |
| Papers from China affiliated with<br>hospitals<br>proportion (%) | 147/159<br>(92%) | **68/73**<br>**(93%)** | 40/44<br>(91%) | **28/31**<br>**(90%)** | 82/90<br>(91%) | **48/50**<br>**(96%)** |
| Papers from all other countries affiliated<br>with hospitals<br>proportion (%) | 6/15<br>(40%) | **0/2**<br>**(0%)** | 0/6<br>(0%) | **0/3**<br>**(0%)** | 1/10<br>(10%) | **0/1**<br>**(0%)** |
| Papers with post-publication notices<sup>b</sup><br>proportion (%) | 20/174<br>(12%) | **13/75**<br>**(17%)** | 1/50<br>(2%) | **1/31**<br>**(3%)** | 1/100<br>(1%) | **1/51**<br>**(2%)** |

<sup>a</sup>SGK papers were published until June 2019<sup>b</sup>Post-publication notices include retractions, expressions of concern and corrections

**Table 2:** Cancer types studied in the Single Gene Knockdown (SGK) corpus, where each cancer type corresponds to a single paper

| Gene | Previously reported SGK papers | New SGK papers |
| --- | --- | --- |
| *ADAM8* | **Liver<sup>a</sup>** | Breast, Breast, Colorectal, Gastric, Liver, Lung, Pancreatic |
| *ANXA1* | N/A | **Breast**, Breast, Breast, Esophageal, Leukemia, Liver, Lung, Prostate |
| *EAG1* | **Liposarcoma, Osteosarcoma** | **Brain, Osteosarcoma**, Osteosarcoma, **Ovarian**, Sarcoma |
| *GPR137* | **Bladder, Brain, <u>Colorectal</u><sup>b</sup>, Pancreatic** | Brain, **Gastric, Leukemia**, Liver, Osteosarcoma, **Ovarian**, Prostate |
| *ICT1* | **<u>Brain</u>** | Breast, Gastric, Leukemia, Lung, **Lymphoma, Prostate** |
| *KLF8* | **Osteosarcoma** | **Bladder, Brain**, Brain, Brain, Breast, Colorectal, Colorectal, **Gastric, Gastric**, Gastric, Gastric, Liver, **Liver, Nasopharyngeal, Oral**, Ovarian, Pancreatic, **Renal** |
| *MACC1* | **Ovarian** | **Bladder**, Brain, **Cervical**, Cervical, Colorectal, Colorectal, **Esophageal**, Esophageal, Gallbladder, Gastric, Liver, Liver, Lung, **Oral, Oral**, Nasopharyngeal, **Ovarian**, Ovarian, **Skin** |
| *MYO6* | **<u>Brain</u>, Colorectal, Liver, Lung** | **Breast**, Gastric, Oral, **Prostate** |
| *NOB1* | Brain, **Breast**, Colorectal, Liver, **Osteosarcoma**, Ovarian, Prostate | Laryngeal, **Lung, Lung**, Oral, Osteosarcoma, Renal, Thyroid, Thyroid |
| *PP4R1* | **<u>Breast, Liver</u>** | **Lung** |
| *PP5* | **Colorectal, Ovarian** | Bladder, **Brain, Leukemia**, Liver, Osteosarcoma, Pancreatic, Prostate |
| *PPM1D* | Bladder, Lung | Brain, Brain, Breast, Breast, Liver, **Pancreatic** |
| *RPS15A* | **Brain, Lung** | Brain, **Gastric, Leukemia**, Liver, Lung, **Osteosarcoma, Renal, Thyroid** |
| *TCTN1* | **<u>Brain</u>**, Brain, **Pancreatic** | Brain, Colorectal, Gastric, Thyroid |
| *TPD52L2* | **Brain, <u>Breast</u>, <u>Gastric</u>, <u>Liver</u>, <u>Oral</u>** | Brain |
| *USP39* | **<u>Liver</u>, Thyroid** | **Breast, Colorectal**, Colorectal, **Gastric, Liver, Liver**, Lung, **Oral**, Osteosarcoma, Renal, Skin |
| *ZFX* | **Brain, Breast** | Brain, Brain, Gallbladder, Laryngeal, Leukemia, Lung, Lung, Oral, Oral, **Osteosarcoma**, Pancreatic, Prostate, **Renal** |

<sup>a</sup>Cancer types shown in bold correspond to problematic papers with wrongly identified nucleotide sequence(s)
<sup>b</sup>Underlined cancer types correspond to papers that have been retracted or assigned an expression of concern

S&B screening (24) flagged 144/174 (83%) SGK papers for further analysis (Fig 1). Manual verification of the identities of all nucleotide sequences in flagged papers (see Methods) confirmed that 75/174 (43%) SGK papers included 1-8 wrongly identified sequences/paper (Table 1, S1 Data). The 75 problematic SGK papers analysed 24 human cancer types, most frequently brain cancer, where 1-9 problematic SGK papers were identified per queried gene (Table 2). Whereas 31/75 (41%) problematic SGK papers have been reported in earlier studies (12, 13), the remaining 45 SGK papers have not been previously analysed (Table 2). The 75 problematic SGK papers were published across 42 journals, where Spandidos Publications published the highest proportion (20/75, 27%) (S1 Data). Problematic SGK papers described 115 wrongly identified sequences (Table 3), where half of these sequences (57/115) targeted a gene or genomic sequence other than the claimed target, followed by incorrect “non-targeting” reagents (44/115, 38%) (Fig 2, Table 3). The 71 incorrect targeting sequences were claimed to interrogate 20 protein-coding genes (S1 Table). Most (67/115, 58%) incorrect sequences recurred across at least 2 SGK papers (Fig 3, S1 Table), where the most frequent incorrect reagent was a previously described “non-targeting” shRNA that is predicted to target *TPD52L2* (11–13). This shRNA or highly similar variants were employed as “non-targeting” controls in 41% (31/75) problematic SGK papers (S1 Table).

**Fig 2.**
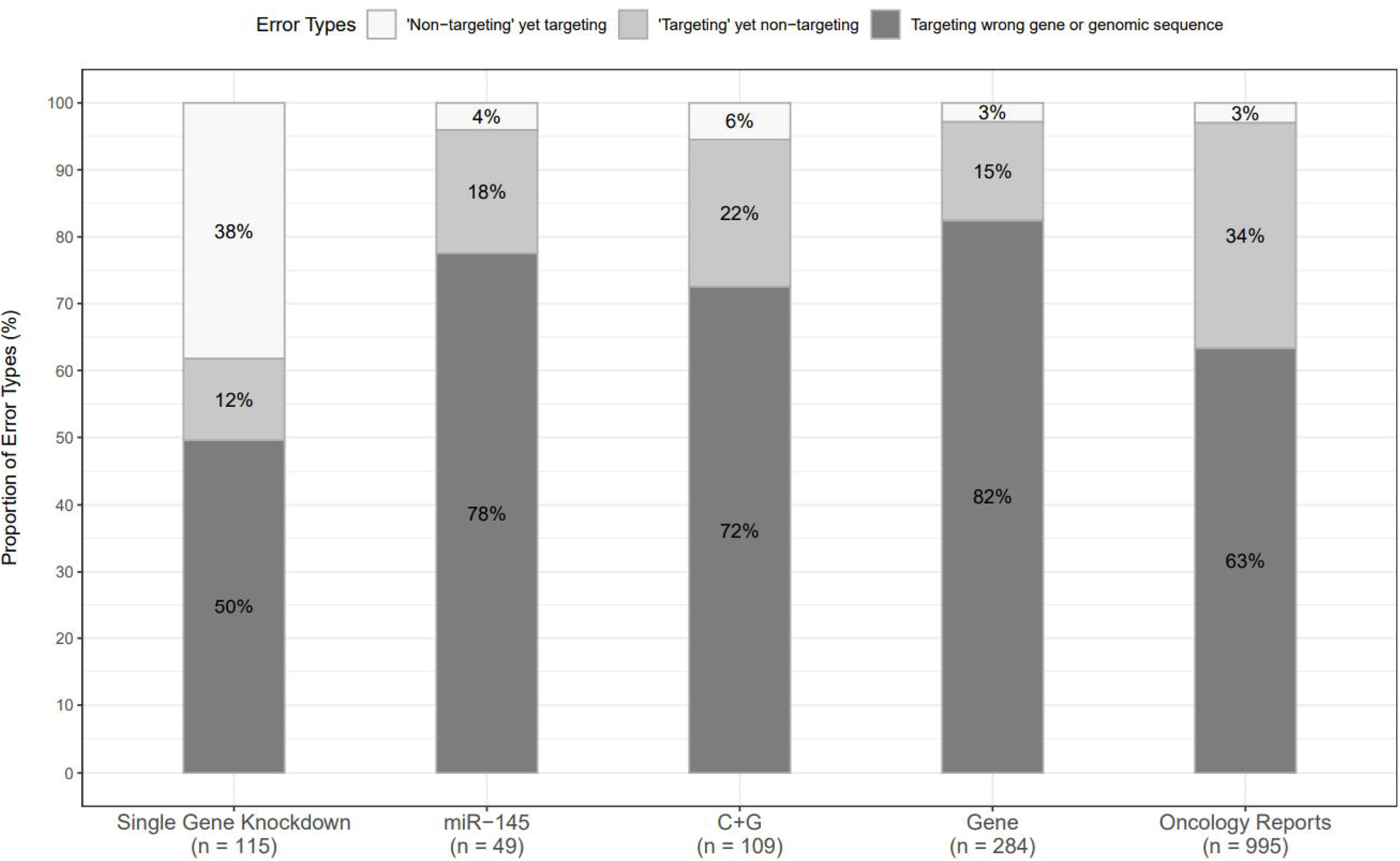
Percentages of sequence identity error types in each corpus. Percentages of wrongly identified nucleotide sequence reagents that correspond to the 3 identity error types (Y axis) in each corpus (X axis). Percentages corresponding to each error type are indicated, rounded to the nearest single digit. The numbers of incorrect sequences in each corpus are shown below the X axis.

**Fig 3.**
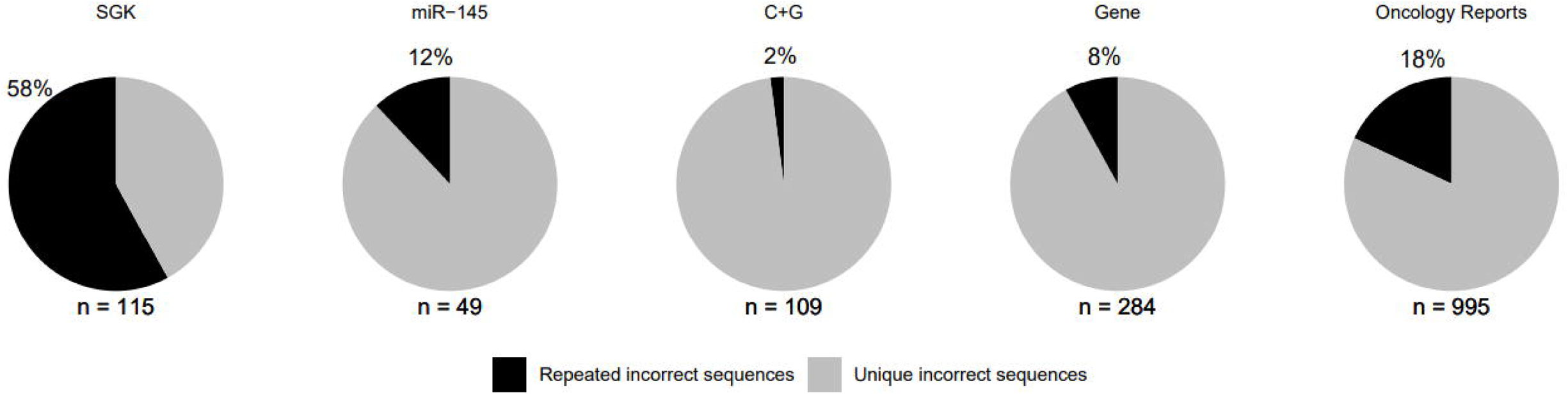
Percentages of wrongly identified nucleotide sequences that were either unique or repeated within each corpus. Percentages of wrongly identified sequences that were identified at least twice in any single corpus (black) are shown above each image, rounded to the nearest single digit. All other wrongly identified sequences were unique in the indicated corpus (grey). Numbers of wrongly identified sequences identified in each corpus are shown below each image.

**Table 3:**
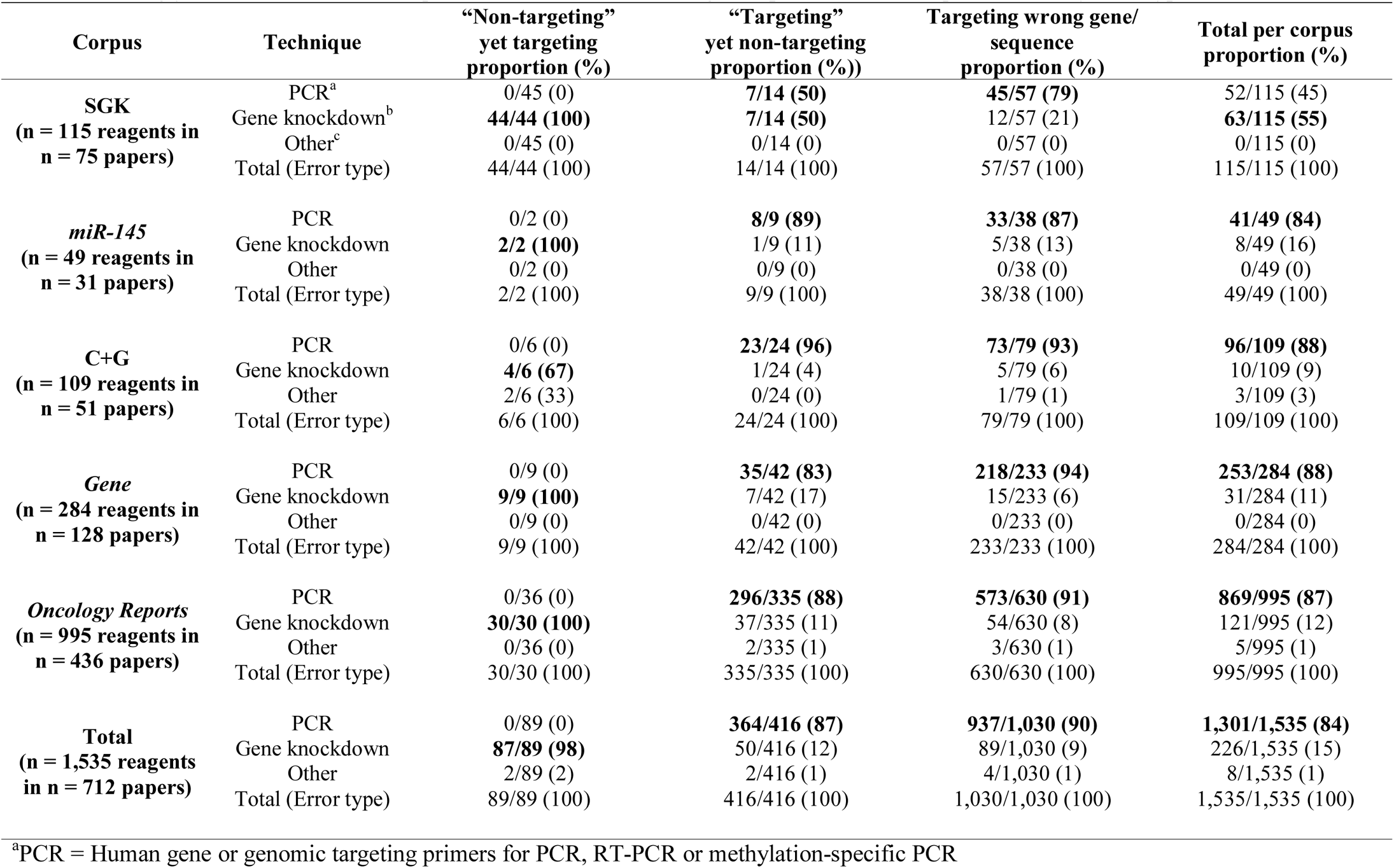
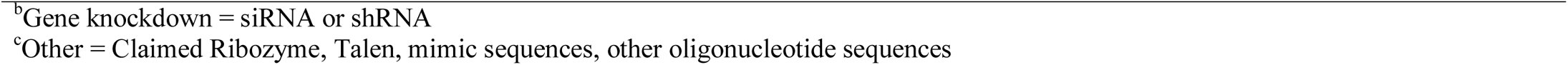
Wrongly identified nucleotide sequences summarized according to experimental technique and identity error type

| Corpus | Technique | “Non-targeting”<br>yet targeting<br>proportion (%) | “Targeting”<br>yet non-targeting<br>proportion (%) | Targeting wrong gene/<br>sequence<br>proportion (%) | Total per corpus<br>proportion (%) |
| --- | --- | --- | --- | --- | --- |
| **SGK**<br>(n = 115 reagents in<br>n = 75 papers) | PCR<sup>a</sup> | 0/45 (0) | **7/14 (50)** | **45/57 (79)** | 52/115 (45) |
|  | Gene knockdown<sup>b</sup> | **44/44 (100)** | **7/14 (50)** | 12/57 (21) | **63/115 (55)** |
|  | Other<sup>c</sup> | 0/45 (0) | 0/14 (0) | 0/57 (0) | 0/115 (0) |
|  | Total (Error type) | 44/44 (100) | 14/14 (100) | 57/57 (100) | 115/115 (100) |
| **miR-145**<br>(n = 49 reagents in<br>n = 31 papers) | PCR | 0/2 (0) | **8/9 (89)** | **33/38 (87)** | **41/49 (84)** |
|  | Gene knockdown | **2/2 (100)** | 1/9 (11) | 5/38 (13) | 8/49 (16) |
|  | Other | 0/2 (0) | 0/9 (0) | 0/38 (0) | 0/49 (0) |
|  | Total (Error type) | 2/2 (100) | 9/9 (100) | 38/38 (100) | 49/49 (100) |
| **C+G**<br>(n = 109 reagents in<br>n = 51 papers) | PCR | 0/6 (0) | **23/24 (96)** | **73/79 (93)** | **96/109 (88)** |
|  | Gene knockdown | **4/6 (67)** | 1/24 (4) | 5/79 (6) | 10/109 (9) |
|  | Other | 2/6 (33) | 0/24 (0) | 1/79 (1) | 3/109 (3) |
|  | Total (Error type) | 6/6 (100) | 24/24 (100) | 79/79 (100) | 109/109 (100) |
| **Gene**<br>(n = 284 reagents in<br>n = 128 papers) | PCR | 0/9 (0) | **35/42 (83)** | **218/233 (94)** | **253/284 (88)** |
|  | Gene knockdown | **9/9 (100)** | 7/42 (17) | 15/233 (6) | 31/284 (11) |
|  | Other | 0/9 (0) | 0/42 (0) | 0/233 (0) | 0/284 (0) |
|  | Total (Error type) | 9/9 (100) | 42/42 (100) | 233/233 (100) | 284/284 (100) |
| **Oncology Reports**<br>(n = 995 reagents in<br>n = 436 papers) | PCR | 0/36 (0) | **296/335 (88)** | **573/630 (91)** | **869/995 (87)** |
|  | Gene knockdown | **30/30 (100)** | 37/335 (11) | 54/630 (8) | 121/995 (12) |
|  | Other | 0/36 (0) | 2/335 (1) | 3/630 (1) | 5/995 (1) |
|  | Total (Error type) | 30/30 (100) | 335/335 (100) | 630/630 (100) | 995/995 (100) |
| **Total**<br>(n = 1,535 reagents<br>in n = 712 papers) | PCR | 0/89 (0) | **364/416 (87)** | **937/1,030 (90)** | **1,301/1,535 (84)** |
|  | Gene knockdown | **87/89 (98)** | 50/416 (12) | 89/1,030 (9) | 226/1,535 (15) |
|  | Other | 2/89 (2) | 2/416 (1) | 4/1,030 (1) | 8/1,535 (1) |
|  | Total (Error type) | 89/89 (100) | 416/416 (100) | 1,030/1,030 (100) | 1,535/1,535 (100) |

<sup>a</sup>PCR = Human gene or genomic targeting primers for PCR, RT-PCR or methylation-specific PCR
<sup>b</sup>Gene knockdown = siRNA or shRNA
<sup>c</sup>Other = Claimed Ribozyme, Talen, mimic sequences, other oligonucleotide sequences

#### miR-145 corpus

PubMed similarity searches employing individual SGK papers identified numerous papers that analysed the functions of different human miR’s in cancer cell lines. We selected *miR-145* from many possible candidates to define a single miR corpus to be screened by S&B. Papers that focussed upon *miR-145* were identified using PubMed similarity searches of index papers (12, 13) and keyword searches of the Google Scholar database (see Methods). A total of 163 *miR-145* papers were then screened by S&B to flag 50 *miR-145* papers for further analysis (Fig 1). The 50 flagged *miR-145* papers were published between 2009-2019 across 35 journals (Table 1), and examined 18 human cancer types, where a single cancer type was analysed in each paper (S2 Data). All flagged papers examined *miR-145* in combination with 1-5 other human gene(s) that were named in publication titles, with a minority of titles (5/50, 10%) also naming a single drug (S2 Data). Most (44/50, 88%) flagged *miR-145* papers were published by authors from China, where most papers (40/44, 91%) were also affiliated with hospitals (Table 1, S2 Data). The 6 *miR-145* papers from 5 other countries were affiliated with institutions other than hospitals (Table 1, S2 Data).

Manual verification of S&B results revealed that most (31/50, 62%) flagged *miR-145* papers described at least one wrongly identified sequence, with a median of one (range 1-5) incorrect sequence/paper (Table 1, S2 Data). The 31 problematic *miR-145* papers were published from 2009-2019 across 25 journals, with the highest proportion published by Wiley (S2 Data). The 31 problematic *miR-145* papers analysed 12 human cancer types, most frequently colorectal or lung cancer, and described 49 wrongly identified sequences, most of which (38/49, 78%) targeted a different gene or target from that claimed (Fig 2, Table 3). The 47 incorrect targeting sequences (Table 3) were claimed to interrogate 13 protein-coding and 4 non-coding RNA’s (ncRNA’s) (S1 Table). In contrast to SGK papers, most incorrect sequences in *miR-145* papers were employed as (RT)-PCR primers (Table 3) and were identified only once within the corpus (Fig 3). All problematic *miR-145* papers were published by authors from China, where almost all papers (29/31, 94%) were affiliated with hospitals (Table 1, S2 Data).

#### Cisplatin and Gemcitabine (C+G) corpus

We noted examples of SGK and *miR-145* papers that analysed the effects of drugs in cancer cell lines (S1 Data, S2 Data). PubMed similarity searches of index papers (12, 13) combined with keyword searches of the Google Scholar database were therefore employed to identify papers that described either cisplatin or gemcitabine treatment of human cancer cell lines and/or cancer patients (see Methods) (Fig 1). A total of 258 papers were screened by S&B to flag 100 papers (n=50 papers for each drug) for further analysis as a combined cisplatin + gemcitabine (C+G) corpus (Fig 1). The 100 flagged C+G papers were published between 2008-2019 across 48 journals (Table 1) and referred to a median of 2 (range 0-4) human genes across their titles (S3 Data). The 100 flagged C+G papers examined 13 human cancer types, where most (96/100, 96%) examined a single cancer type, typically pancreatic (35/100, 35%) or lung (22/100, 22%) cancer (S3 Data), reflecting the clinical use of cisplatin and gemcitabine (27, 28). Most (90/100, 90%) C+G papers were published by authors from China, where most (82/90, 91%) were also affiliated with hospitals (Table 1, S3 Data). In contrast, 1/10 C+G papers from 8 other countries was hospital-affiliated (Table 1, S3 Data).

Approximately half (51/100, 51%) the flagged C+G papers were found to include a median of 2 (range 1-8) wrongly identified sequences/paper (Table 1). The 51 problematic C+G papers were published between 2009-2019 across 31 journals (Table 1), where Springer Nature published the highest proportion (15/51, 29%), followed by Elsevier (12/51, 24%) (S3 Data). The 51 problematic C+G papers examined 13 human cancer types, most frequently pancreatic cancer, and described 109 wrongly identified nucleotide sequences, most of which (79/109, 72%) targeted a gene or genomic sequence other than the claimed target (Fig 2, Table 3, S1 Table). The 103 incorrect targeting sequences (Table 3) were claimed to interrogate 31 protein-coding genes and 16 ncRNA’s (S1 Table). As in *miR-145* papers, most incorrect sequences in problematic C+G papers represented (RT)-PCR primers (Table 3) and were identified once within the corpus (Fig 3). Almost all (50/51, 98%) problematic C+G papers were published by authors from China, where almost all (48/50, 96%) were affiliated with hospitals (Table 1, S3 Data).

### Analysis of Gene and Oncology Reports corpora

As papers in targeted corpora were selected using known features of human research papers with incorrect nucleotide sequences, we complemented these analyses by screening original research papers in the journals *Gene* and *Oncology Reports*. These journals were selected as examples of journals that have published papers with incorrect nucleotide sequences (11–13), where *Oncology Reports* also published the highest number of problematic SGK papers (S1 Data). S&B was employed to screen all original articles published in *Gene* from 2007-2018, and all open-access articles published in *Oncology Reports* from 2014-2018 (Table 4).

**Table 4:** Summary of features of *Gene* and *Oncology Reports* journals and problematic papers

| **Feature** | ***Gene*** | ***Oncology Reports*** |
| --- | --- | --- |
| Publication years screened by Seek & Blastn | 2007-2018 | 2014-2018 |
| Journal Impact Factor<br>(range during years screened) | 2.082-2.871 | 2.301-3.041 |
| Flagged/ screened papers<br>proportion (%) | 742/7,399 (10%) | 1,709/3,778 (45%) |
| Problematic/ flagged papers<br>proportion (%) | 128/742 (17%) | 436/1,709 (26%) |
| Incorrect sequences/ problematic paper<br>median (range) | 2 (1-36) | 2 (1-15) |
| Problematic papers from China<br>proportion (%) | 69/128 (54%) | 393/436 (90%) |
| Problematic papers from all other countries<br>proportion (%) | 59/128 (46%) | 43/436 (10%) |
| Problematic papers from China affiliated with<br>hospitals<br>proportion (%) | 54/69 (78%) | 342/393 (87%) |
| Problematic papers from all other countries<br>affiliated with hospitals<br>proportion (%) | 5/59 (9%) | 5/43 (12%) |
| Retracted or corrected problematic papers<br>proportion (%) | 2/128 (2%) | 2/436 (0.5%) |

Screening 7,399 original *Gene* articles from 2007-2018 flagged 742 (10%) papers for further analysis (Fig 1) (see Methods). Manual verification of S&B outputs found that 17% (128/742) flagged papers described a median of two (range 1-36) wrongly identified sequences/paper (Table 4, S4 Data). These 128 problematic papers referred to 186 human genes (n=146 protein-coding, n=40 ncRNA’s) across their publication titles (S4 Data). Approximately half (65/128, 51%) the problematic *Gene* papers analysed gene function in research contexts other than human cancer, most frequently by examining gene polymorphisms in patient cohorts (16/65, 25%) (S4 Data). The remaining 60 papers analysed 17 different human cancer types, most frequently lung cancer (12/60, 20%) (S4 Data). A minority of problematic *Gene* papers (7/128, 5%) referred to drugs within their titles. Manual verification of over 5,200 sequences highlighted 284 wrongly identified sequences across the 128 problematic *Gene* papers. Almost all (275/284, 97%) incorrect sequences represented targeting reagents (Fig 2, Table 3) for the analysis of 92 protein-coding genes and 24 ncRNA’s (S1 Table). Most (261/279, 92%) incorrect sequences were described once within the *Gene* corpus (Fig 3).

As *Oncology Reports* published many more papers per year than *Gene* from 2007-2018, we employed S&B to screen open-access *Oncology Reports* articles from 2014-2018 (n=3,778 papers, 99% *Oncology Reports* articles) (Fig 1, Table 4). Almost half (1,709/3,778, 45%) screened papers were flagged for further analysis (Fig 1), and over one quarter (436/1,709, 26%) of flagged papers were confirmed to describe a median of 2 (range 1-15) wrongly identified sequences/paper (Table 4, S5 Data). Almost all (432/436, 99%) problematic *Oncology Reports* papers studied gene function in human cancer, most frequently lung (54/432, 13%) or liver cancer (46/432, 11%). A subset (51/432, 12%) of problematic papers referred to 42 different drugs across their titles, most frequently cisplatin or 5-fluorouracil (S5 Data). Manual verification of over 5,100 sequence identities confirmed 995 wrongly identified sequences (S1 Table). Almost all (965/995, 97%) incorrect sequences represented targeting reagents (Fig 2, Table 3) for the analysis of 262 protein-coding genes and 86 ncRNA’s (S1 Table). Most (816/965, 85%) incorrect sequences were described once across the *Oncology Reports* corpus (Fig 3).

#### Geographic, institutional and temporal distributions of problematic Gene and Oncology Reports papers

The 128 problematic *Gene* papers were authored by teams from 19 countries (S1A Fig) (see Methods). Just over half (69/128, 54%) problematic *Gene* papers were authored by teams from China (Table 4, Fig 4, S1B Fig), followed by India (10/128, 8%) and Iran (9/128, 7%) (S1A Fig). Similar results were obtained for the 95 problematic *Gene* papers from 2014-2018, where teams from China authored 66% (63/95) papers (Fig 4). A significantly greater proportion of problematic *Gene* papers from China were affiliated with hospitals (54/69, 78%), compared with papers from other countries (5/59, 8%) (Fisher’s exact test, p<0.001, n=128 papers) (Fig 4). This difference was also noted for problematic *Gene* papers from 2014-2018 (Fisher’s exact test, p<0.001, n=98 papers) (Fig 4).

**Fig 4.**
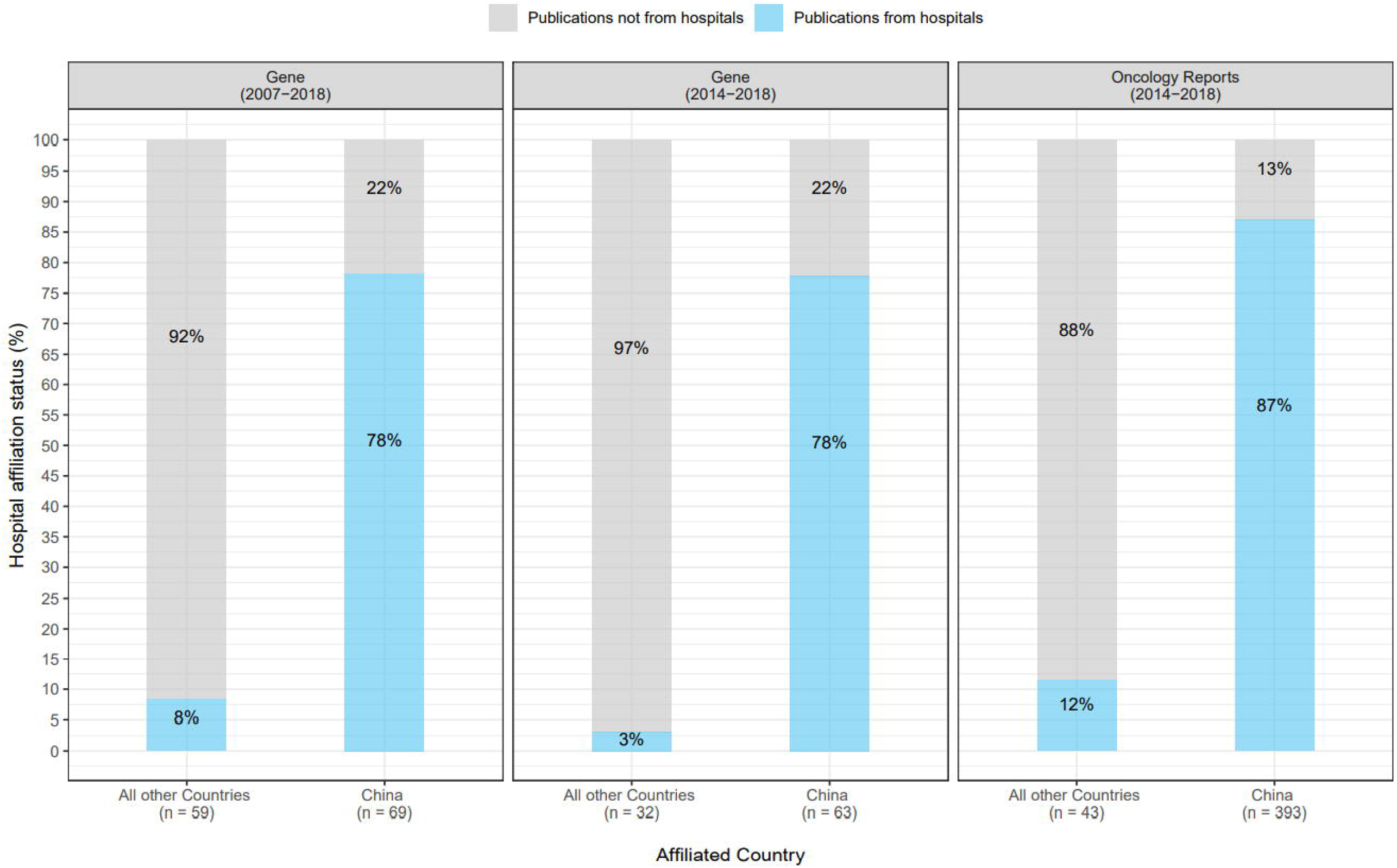
Percentages of problematic *Gene* and *Oncology Reports* papers according to hospital affiliation status and country of origin. Percentages of problematic *Gene* and *Oncology Reports* papers according to hospital affiliation status (Y axis) from either China or all other countries (X axis). The journal and relevant date ranges of problematic papers are shown above each panel. Problematic papers that were (not) affiliated with hospitals are shown in blue (grey), respectively. Percentages shown have been rounded to the nearest single digit. Numbers of problematic papers from China or all other countries are indicated below the X axis. For the comparisons shown in each panel, significantly higher proportions of problematic papers from China were affiliated with hospitals versus problematic papers from other countries (Fisher’s Exact text, p<0.001).

In contrast to the broader geographic distribution of problematic *Gene* papers, the 436 problematic *Oncology Reports* papers were authored by teams from just 13 countries (S2A Fig). Most (393/436, 90%) problematic *Oncology Reports* papers were authored by teams from China (Table 4, Fig 4), followed by much smaller proportions from South Korea (14/436, 3%) and Japan (12/436, 3%) (S2A Fig). As noted for problematic *Gene* papers, a significantly greater proportion of problematic *Oncology Reports* papers from China were affiliated with hospitals (342/393, 87%) compared with papers from other countries (5/43, 12%) (Fisher’s exact test, p<0.001, n=436 papers) (Fig 4).

We considered the distributions of problematic *Gene* and *Oncology Reports* papers according to year of publication, country of origin, and affiliated institution type (Fig 5, S1 Fig, S2 Fig). Problematic *Gene* papers were infrequent from 2007-2011 (1-4 papers/year), rising to 8-38 papers/year from 2012-2018, where the highest number of problematic papers was identified in 2018 (Fig 5). These numbers correspond to 1.0%-4.2% of all original *Gene* papers published per year from 2012-2018. Papers from China represented the majority of problematic *Gene* papers from 2015-2018 (Fig 5), where most papers were also affiliated with hospitals (S1B Fig). Compared with *Gene*, *Oncology Reports* published higher numbers of problematic papers per year, corresponding to 8.3-12.6% original *Oncology Reports* papers from 2014-2018 (Fig 5). Across all 5 years, most (87-93%) problematic *Oncology Reports* papers were authored by teams from China, corresponding to 11% original *Oncology Reports* articles in 2015-2017 (Fig 5), most of which were also affiliated with hospitals (S2B Fig).

**Fig 5.**
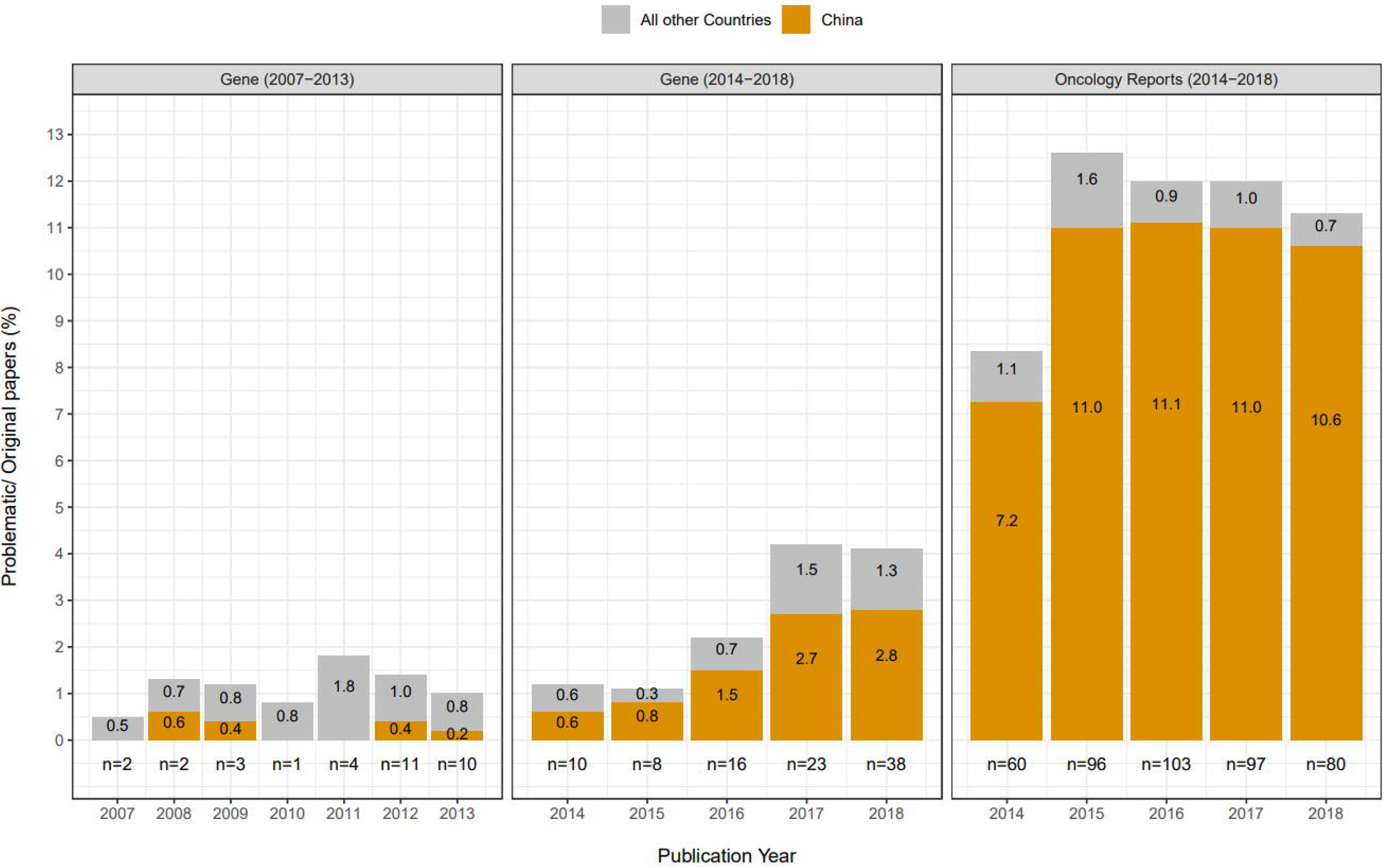
Percentages and numbers of problematic *Gene* and *Oncology Reports* papers per year. Percentages of all *Gene* or *Oncology Reports* papers that were found to be problematic (Y axis) per publication year (X axis). The journal and relevant publication year ranges are shown above each panel. Problematic papers from China or all other countries are shown in orange or grey, respectively. Percentages shown are rounded to one decimal place. Total numbers of problematic papers per year are shown below each graph.

### Analysis of all problematic human gene function papers

After adjusting for 9 duplicate papers across the 5 corpora, we identified 712 problematic papers with wrongly identified sequences (Fig 1) that were published by 78 journals and 31 publishers (S6 Data). The 712 problematic papers included 1,535 wrongly identified sequences, most of which were (RT-)PCR reagents (1,301/1,535, 85%), followed by si/shRNA’s (226/1,535, 15%) (Table 3).

As most incorrect reagents represented (RT-)PCR primers which are employed as paired reagents, we considered the verified identities of primer pairs that were found to include at least one wrongly identified primer (Fig 6). Problematic papers frequently paired one (RT-)PCR primer that targeted the claimed gene with a primer that was predicted to target a different gene (n=237 papers), or to be non-targeting in human (n=118 papers) (Fig 6). Many problematic papers (n=192) described primer pairs that were predicted to target the same incorrect gene (Fig 6). Problematic papers also combined one non-targeting primer with another that was predicted to target an incorrect gene (n=70 papers), two non-targeting primers (n=63 papers), and/or primers that were predicted to target two different incorrect genes (n=42 papers) (Fig 6). Notably, 21% (276/1,301) incorrect (RT-)PCR primers were predicted to target an orthologue of the claimed gene, typically in rat or mouse (S1 Table).

**Fig 6.**
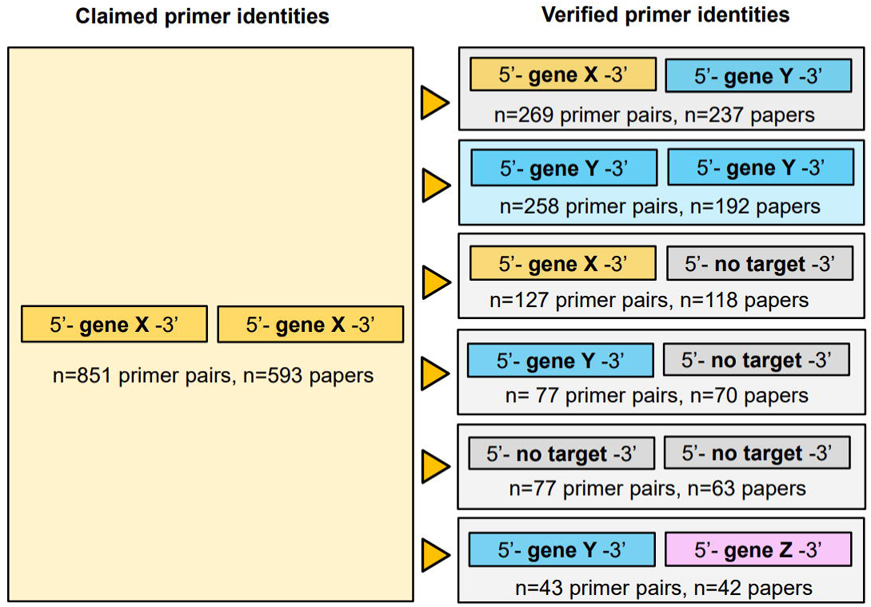
Summary of (RT-)PCR primer pairings that involved at least one wrongly identified primer. For n=851 primer pairs that were claimed to target particular genes/sequences (gene X) (left panel), one or both primers were predicted to be incorrect (right panel), either by targeting unrelated genes or sequences (gene Y or gene Z), or by having no predicted human target (no target). Numbers of primer pairs and affected papers are indicated below each incorrect primer pair category. Some problematic papers described more than one (category of) incorrect primer pairing. Left- or right-hand primers are not intended to indicate forward or reverse primer orientations.

#### Bibliometric analysis of human genes analysed in problematic papers

Almost all (1,442/1,535, 94%) incorrect sequences represented targeting reagents that were claimed to target 365 protein-coding genes and 120 ncRNA’s (S1 Table). The remaining 88 sequences represented incorrect “non-targeting” sequences that were instead predicted to target 35 genes, most of which (28/35, 80%) were protein-coding genes.

To count the numbers of papers in PubMed that are associated with protein-coding genes in n=709 problematic papers (Fig 7), we used the gene2pubmed service of the National Center for Biotechnology Information (29), restricting these analyses to protein-coding gene identifiers that mapped to official gene names. Primary protein-coding genes, which represented the first-listed genes in publication titles or abstracts, tended to be associated with more papers in PubMed than a randomly chosen human protein-coding gene (median publication numbers: 167 vs 31, *P* < 10^−109^, two-sided Mann-Whitney U test) (S3A Fig). Only two genes that were the primary focus of least two problematic papers (*TCTN1* and *GPR137)* (Table 2) have appeared in fewer publications in PubMed than a randomly chosen human protein-coding gene (Fig 7A). We repeated these analyses to examine the protein-coding genes that were claimed as targets by wrongly identified reagents. Again, most wrongly identified target genes have appeared in more papers than a randomly chosen protein-coding gene (median publication numbers: 238 vs 31, *P* < 10^−94^, two-sided Mann-Whitney U test) (S3B Fig). The most frequent wrongly claimed gene targets were *GAPDH* and *ACTB* (Fig 7B), reflecting their widespread use as RT-PCR control genes. In summary, these analyses demonstrate that problematic papers can focus upon and/or employ reagents that are wrongly claimed to target highly-investigated human genes such as *BCL2*, *EGFR*, *PTEN*, *STAT3*, and *CCND1* (Fig 7).

**Fig 7.**
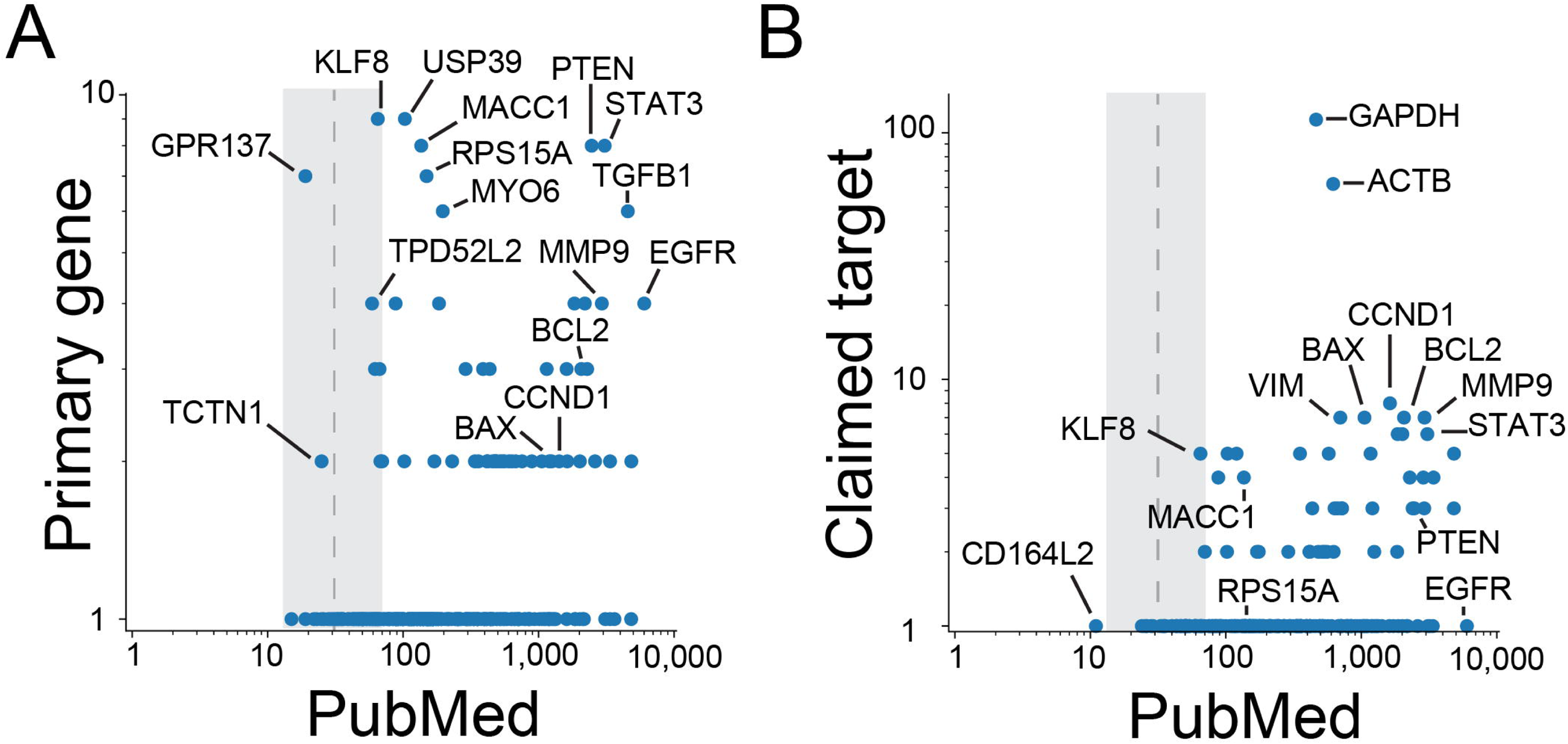
Numbers of past research papers that have studied human protein-coding genes in problematic papers. Numbers (log base 10) of problematic papers (Y axis) versus past research papers (X axis) for A: primary protein-coding genes in problematic papers and B: claimed protein-coding gene targets of wrongly identified reagents. Vertical dashed lines indicate the median number of research papers for protein-coding genes, with the associated interquartile range shown in grey. Subsets of protein-coding genes are highlighted in each panel.

#### Post-publication correction, citation and curation of problematic papers

We considered whether any problematic papers have been the subject of post-publication notices, such as retractions, expressions of concern or corrections (11). Only 2% (11/712) of problematic papers have been retracted, where most (8/11) retraction notices did not refer to wrongly identified sequence(s), and 3 problematic papers have been subject to expressions of concern (S2 Table). Although we excluded papers in which incorrect sequences had been subsequently corrected (see Methods), we noted 5 corrections to problematic papers that addressed issues other than incorrect sequences (S2 Table). Almost all (693/712, 97%) problematic papers therefore remain uncorrected within the literature.

We then considered how problematic papers have been curated within gene knowledgebases and cited within the literature. Between 1-207 problematic papers were found within 5 gene knowledgebases that rely upon text mining (30–34), where knowledgebases of miR functions contained the most problematic papers (S3 Table). In March 2021, the 712 problematic papers had been cited 17,183 times according to Google Scholar. Subsets of problematic C+G, *Gene* and *Oncology Reports* papers have also been cited by one or more clinical trials (Fig 8A). Given expected publication delays between pre-clinical and clinical research, we extended these data by considering the approximate potential to translate (APT) for problematic papers (35) according to publication corpus (Fig 8B). The APT metric uses the combination of concepts contained within a paper to infer the probability that the paper will be cited by future clinical trials or guidelines (35). The average APT for problematic papers in the 5 corpora ranged from 15-35% (Fig 8B), indicating that 15-35% of problematic papers in each corpus resemble papers that will be cited by clinical research. Without timely interventions, or without heuristics not captured by the models underlying the APT (35), around one quarter of problematic papers are likely to directly misinform or distract the future development of therapies against human disease.

**Fig 8.**
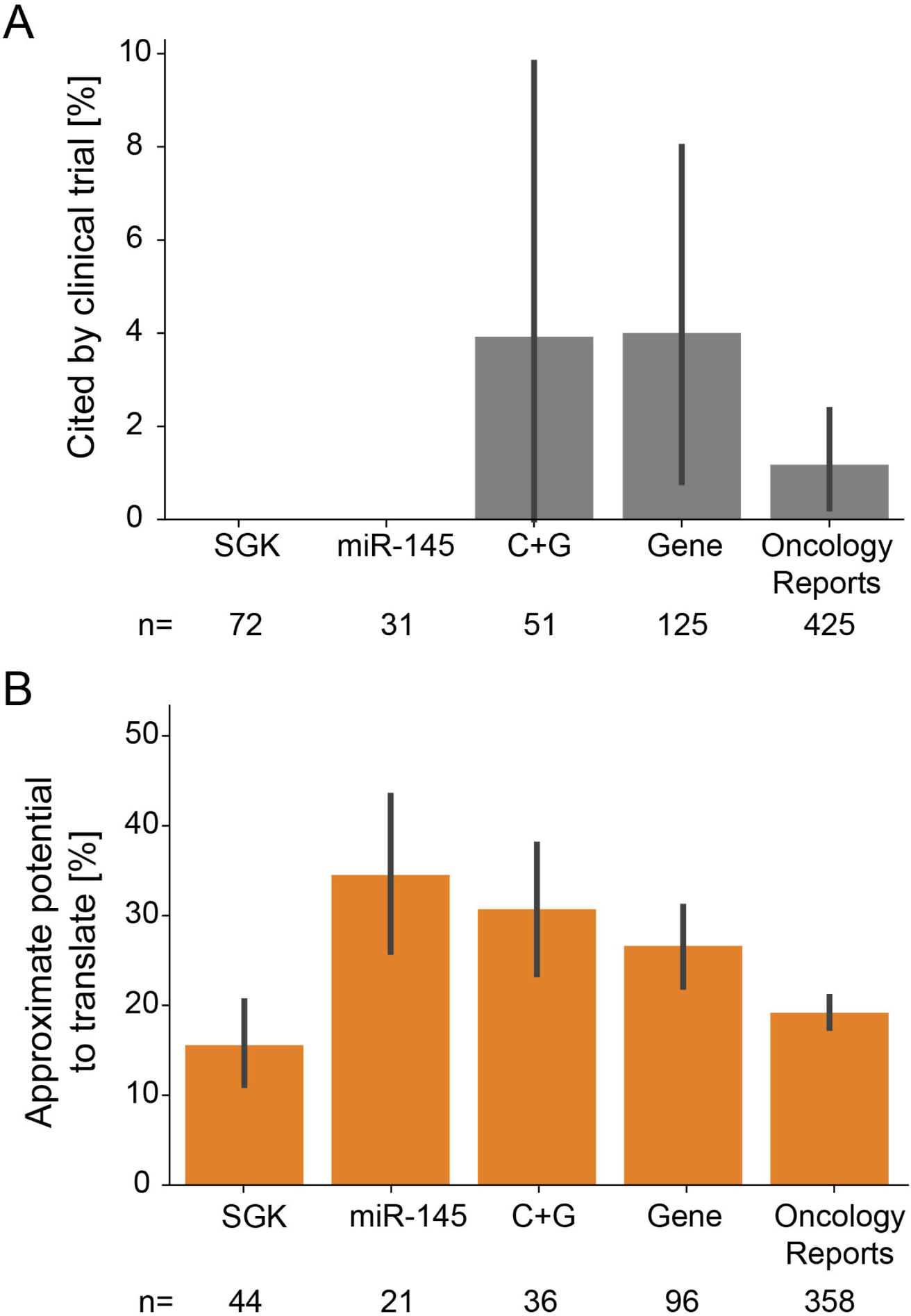
Clinical trial citations and approximate potential to translate for problematic papers. A: Percentages of problematic papers that are cited at least once according to the NIH Open Citation Collection (Y axis), according to publication corpus (X axis). Error bars indicate 95% confidence intervals of bootstrapped estimates of percentages. Numbers of problematic papers with at least one clinical citation are shown below the X axis for each corpus. B: Average approximate potential to translate for problematic papers (Y axis) according to publication corpus (X axis). Error bars indicate bootstrapped 95% confidence intervals. Numbers of problematic papers for which the approximate potential to translate was computed by iCite are shown below the X axis for each corpus.

## Discussion

Experimental analyses of gene function require nucleotide sequence reagent identities to precisely match their published descriptions. Wrongly identified nucleotide sequence reagents therefore represent a threat to the continuum of genetics research, from population-based genomic sequencing to pre-clinical functional analyses, and the translation of these results to patients. While there have been previous reports of wrongly identified PCR and gene knockdown reagents in single papers or small cohorts (11–21), the present study is the first to systematically fact-check the identities of nucleotide sequences in over 3,400 papers. Our supported application of S&B (12, 24, 26) for both targeted corpora and journal screening identified 712 papers in 78 journals that describe 1-36 wrongly identified nucleotide sequences/paper. These 712 papers described over 1,500 wrongly identified sequences, most of which were claimed targeting reagents for the study of over 480 human genes. The sheer number of problematic papers and incorrect reagents that we have uncovered from screening a very small fraction of the human gene function literature predicts a problem of alarming proportions that requires urgent and co-ordinated action.

Before outlining the possible significance of our results, we must recognise the limitations of the present study. Firstly, we recognise that by verifying the identities of over 13,700 sequences, we are likely to have committed some errors of our own. The most challenging incorrect reagents that we encountered are claimed human targeting sequences that appear to have no human target. As blastn is indicated to have a very low but measurable false-negative rate (36, 37), we acknowledge that a small fraction of what appear to be non-targeting reagents may in fact be targeting reagents, as claimed. Nonetheless, as we have taken extensive steps to verify the identities of individual nucleotide sequences (see Methods), we reasonably expect that false-positives at both the sequence and publication level will be rare within our dataset. Many papers also described at least two wrongly identified nucleotide sequences, building in some protection against false-positives at the publication level.

We also recognise that we have screened three targeted corpora, of which two (the *miR-145* and C+G corpora) did not include all available papers. We therefore recognise that the rates of problematic papers within these corpora may not reflect the rates of problematic papers in other targeted corpora or in the wider literature. Nonetheless, the relative proportions of nucleotide sequence error types in problematic *miR-145* and C+G papers were similar to those identified by screening papers in both *Gene* and *Oncology Reports* (Fig 2). This indicates that features of problematic papers identified within the targeted corpora may extend to papers elsewhere, particularly as problematic *miR-145* and C+G papers were published across 47 journals.

Finally, we recognize that by only verifying the S&B outputs for a minority of papers published in *Gene* and *Oncology Reports*, we will not have described all papers with wrongly identified nucleotide sequences in these journals. Indeed, the identified numbers and proportions of problematic papers are very likely to represent underestimates, as these proportions were derived from verifying sequences in just 10% *Gene* papers from 2007-2018 and 45% *Oncology Reports* papers from 2014-2018. As S&B does not analyze papers where most species identifiers correspond to species other than human (26), and we did not verify the identities of sequences that were claimed to target non-human species, we also recognize that our results do not indicate whether wrongly identified nucleotide sequences occur in papers that examine other species. Analyses of targeted corpora identified around reference papers may begin to answer this question.

Despite these limitations, our results paint a worrying picture of the reliability of some sections of the human gene-focussed literature. The consequences of many incorrect gene function papers therefore deserve consideration. As previously discussed, papers that describe incorrect nucleotide sequences could encourage the incorrect selection of genes for further experimentation, possibly at the expense of more productive candidates (8). This could be exacerbated where multiple problematic papers report similar results for the analysis of the same gene (8, 13). For example, series of 3-20 single gene knockdown papers that universally claim that the gene target plays a causal role in 3-11 different cancer types could collectively encourage further research. Many incorrect gene function papers could also lead to the overestimation of knowledge of gene function from text mining approaches (30–34), particularly given the assumed reliability of published experimental results (38). Our results demonstrate that problematic papers are already indexed within gene function knowledge bases (30–34).

As experimental reagents, wrongly identified nucleotide sequences carry the additional risk of being wrongly used in other studies (8, 11). Although siRNA’s and shRNA’s are increasingly purchased from external companies as preformulated reagents, many researchers continue to order custom-made PCR primers. The most frequent incorrect reagent type that we have identified can therefore easily be reused from the literature. Experiments that either attempt to replicate published results associated with incorrect reagents and/or unknowingly reuse incorrect reagents are likely to generate unexpected results that may then remain unpublished. As possible evidence of this, we could only identify one study that reported incorrect sequences based on the results of follow-up experiments (20) that our analyses also identified. Given that over 17,000 citations have been accumulated by the problematic papers that we have identified, it seems inevitable that unreliable gene function papers are already wasting time and resources.

Given the numbers of problematic papers and incorrect reagents that we have identified, we must consider possible explanations for both incorrect nucleotide sequences and the papers that describe them. As nucleotide sequences are prone to being affected by different error types, we believe that wrongly identified sequences represent unintended errors that persist when authors, editors and peer reviewers do not check sequence identities during manuscript preparation or review (8, 11, 12). We recognize that wrongly identified sequences in papers could also reflect some form of research sabotage. However, as workplace sabotage is typically directed towards known individuals (39, 40), this seems an unlikely explanation for wrongly identified sequences across hundreds of gene function papers published by many different authors.

We also recognize that some of the wrongly identified sequences that we have described have almost certainly arisen in the context of genuine research. Nonetheless, some wrongly identified sequences were associated with results that appeared highly implausible. For example, the majority of incorrect (RT-)PCR primer pairs that we identified should have uniformly failed to generate (RT-)PCR products across all templates, and yet appeared to generate results that were consistent with primers targeting the claimed genes. As previously reported (11–13), we also identified papers where the non-targeting si/shRNA was verified to target the gene of interest and yet still generated the expected negative or baseline results. Based on the many discrepancies between verified sequence identities and their associated results, it would appear that many experiments across hundreds of gene function papers were not performed as described.

Although most analyses of experimental errors consider the context of genuine research (41–43), errors can also arise within the context of fraudulent or fabricated research (43, 44). We have previously proposed that incorrect nucleotide sequences could flag that some affected papers are fraudulent (8, 11–13, 45, 46). Due to numerous unexpected similarities between single gene knockdown papers with incorrect sequences, we initially proposed that these papers may reflect the undeclared involvement of organisations such as paper mills (13) which have been alleged to mass-produce fraudulent manuscripts for publication (47). The production of fraudulent manuscripts at scale may involve the use of manuscript templates, leading to unusual degrees of similarity between the resulting publications (8, 13, 46). Papers with incorrect nucleotide sequences showed features consistent with the possible use of manuscript templates, such as similarities in textual and figure organisation, outlier levels of textual similarity, superficial explanations for the analysis of particular genes, and generic experimental approaches regardless of predicted gene function (8, 13, 45, 46). In addition to the use of manuscript templates, we have proposed that producing many gene-focused manuscripts at minimal cost could involve writers with either an incomplete understanding of the experiments that they are describing and/or limited time for quality control (8, 46). These conditions could lead to incorrect nucleotide sequences being a feature of gene function manuscripts from paper mills (8, 46) where incorrect sequences could also be reused across different manuscripts (11–13). The description of errors which are incompatible with reported results and the presence of multiple incorrect sequences across different techniques within single papers could therefore be features of experiments that were never actually performed.

We have proposed that human genes represent attractive publication targets for paper mills, with many under-studied human genes (48–50) that can be targeted in different cancer or disease types that can then be distributed across different authors and journals over many years (8, 45, 46). However, whereas we had previously hypothesised that under-studied genes might be preferentially targeted by paper mills (8), in the present analysis, problematic papers rarely focussed on under-studied human protein-coding genes, and instead either focussed on or employed incorrect reagents that were claimed to target protein-coding genes that had received prior attention. While under-studied genes may present more individual publication opportunities, papers that describe human genes with known functions and significance could carry more editorial and reader interest, increasing the likelihood of problematic manuscripts being accepted for publication and then cited by future research.

Our analyses of both targeted and journal corpora indicate that ncRNA’s may provide a further layer of possibilities for the fabrication of gene-focussed papers. While we recognise that studying genes in different diseases and analysing the functions of ncRNA’s in combination with other genes are features of genuine research, our results suggest that a focus upon ncRNA’s such as miR’s could allow the inclusion of more topic variables within manuscripts from paper mills, such as ncRNA’s and protein-coding genes that are studied across different disease types, with or without drug or natural product treatments. Examining ncRNA’s in combination with other gene(s) could allow larger and more diverse publication series to be created, compared with those that focus on single genes. Furthermore, as ncRNA’s possess largely numeric identifiers that may be more difficult to recognise and recall than alphanumeric gene identifiers, any focus upon ncRNA’s could contribute to large publication series being less visible within the literature. As papers that describe miR functions have also been shown to be highly cited (51), miR’s and other ncRNA’s could represent attractive target genes for paper mills.

In both targeted and journal corpora, most problematic papers were authored by teams from mainland China that were also overwhelmingly affiliated with hospitals. While researchers in different countries may use paper mills to meet publication targets or quotas (49), paper mills have been most widely discussed in the context of academics and medical doctors in China (47, 53–59). Stringent publication requirements may represent a particular challenge for hospital doctors in China, where some hospital doctors have described limited time, training and/or opportunities to undertake research (47, 53–59). Large numbers of human gene function papers with incorrect nucleotide sequences that list hospital affiliations in China could reflect hospital doctors turning to paper mills to meet publication requirements, whereas the contrasting institutional profiles of problematic papers from other countries could highlight different publication pressures elsewhere. In summary, our results combined with previous reports (47, 53–59) indicate that publication pressure upon hospital doctors in China may be exerting measurable effects upon the human gene function literature.

In summary, we are concerned that the sheer number of human genes that are available for analysis, combined with research drivers that favour the continued investigation of genes of known function (48–50), are unwittingly providing an extensive source of topics around which gene function papers can be fraudulently created. Furthermore, since genuine pre-clinical gene research requires specialised expertise, time and material resources (13), the mass production of fraudulent gene function papers could be quicker and cheaper by orders of magnitude (8). Given the number of human genes which can be studied either singly or in combination with other genes and topics such as drugs and analysed across different cancer types or other diseases, combined with unrealistic demands for research productivity (46, 53–59), the publication of fraudulent gene function papers could potentially outstrip the publication of genuine gene function research.

### Future directions

Our results indicate that the problem of incorrect gene function papers requires urgent action. Within the research community, this can take place in several ways. As the validation of nucleotide sequence identities using algorithms such as blastn (12, 25) represents a routine activity for teams that investigate gene function, we hope that our results will encourage researchers to unfailingly check the identities of published nucleotide sequences, both in the context of their own research and during peer review. Researchers encountering wrongly identified sequences can describe these to authors, journal(s) and/or PubPeer (60) using the reporting fields that we have proposed (11). Researchers can also compare the claims of gene-focused papers with those from high-throughput experimental studies (61) and/or predictive algorithms (38, 62). Professional societies can reinforce the importance of reagent verification through conference presentations, education programs, and journal editorials, and can advocate for tangible incentives to encourage further fact-checking of the genetics literature.

Our analysis of only a small proportion of human gene function papers, combined with the discovery that most incorrect nucleotide sequences are unique within screened literature corpora, highlights the need for further screening studies to identify other problematic gene function papers. Journals that published problematic papers in targeted corpora represent possible targets for future screening approaches. As incorrect nucleotide sequences are unlikely to be found in all problematic gene function papers, future research should also combine analyses of nucleotide sequences with other features of concern such as manipulated or recurring experimental images and data (46, 63, 64).

Unfortunately, efforts from the research community alone will not solve the problem that we have described. Similarly, recent changes to researcher assessment (65, 66) will not address problematic papers that have already been published. The ability to alter the published record relies upon the engagement and co-operation of journals and publishers (11). Over the past year, growing numbers of journals have begun to recognize the issue of manuscripts and publications from paper mills (67–77), including papers that analyse gene function and drug treatment in cell lines (71, 74). While the described efforts to screen incoming manuscripts are welcome and should be extended to all journals that publish gene function research, screening incoming manuscripts must be coupled with addressing problematic papers that are already embedded in the literature (71, 74–76). These efforts could be supported by gene function experts who could explain the significance of incorrect nucleotide sequences and/or provide training for editorial staff, particularly as the necessary researcher skills are already widely available. To overcome the protracted timeframes that can be associated with journal investigations of incorrect sequences (11), we have proposed the rapid publication of editorial notes to transparently flag papers with verifiable errors while journal and institutional investigations proceed (46).

### Summary and conclusions

To fully extend the benefits of genomics towards patients and broader populations, it is widely recognised that we must understand the functions of every human gene (1, 2). However, the availability of many human genes for experimental analysis, combined with research drivers that favour the continued investigation of genes of known function (46–49), may unwittingly provide an extensive source of topics around which gene function papers can be fraudulently created. Whereas genuine gene research requires time, expertise, and material resources, the mass production of fraudulent gene function papers by paper mills could be quicker and cheaper by orders of magnitude (8). Given the number of human genes whose functions can analysed singly and/or in combination with other genes and/or drugs across different cancer types or other diseases, combined with acute demands for research productivity that may not always be matched by researcher capacity and training (78), fraudulent gene function papers could unfortunately outstrip the publication of genuine gene function research. Indeed, the possible extent of the problem of unreliable human gene function papers is indicated by the lack of overlap between the problematic papers that we have reported, and other papers of concern reported elsewhere (71, 74, 79). While publishers and journals decide how to address this urgent problem, laboratory scientists, text miners and clinical researchers must approach the human gene function literature with a critical mindset, and carefully evaluate the merits of individual papers before acting upon their results.

## Materials and Methods

### Identification of literature corpora

#### Single Gene Knockdown (SGK) corpus

Single gene knockdown (SGK) papers were identified by combining each of 17 human gene identifiers (*ADAM8, ANXA1, EAG1, GPR137, ICT1, KLF8, MACC1, MYO6, NOB1, PP4R1, PP5, PPM1D, RPS15A, TCTN1, TPD52L2, USP39,* and *ZFX)* with the search string “cancer AND/OR knockdown AND/OR lentivirus” (13), to search PubMed and Google Scholar databases in June 2019 using the “allintext:” function for Google Scholar searches. No publication date ranges, country-specific or journal-based search terms were used to limit search results. Papers were visually inspected to confirm that papers described gene knockdown experiments that targeted one of the 17 human genes in human cancer cell lines.

#### miR-145 corpus

The *miR-145* corpus included papers that analysed human *miR-145* function in human cell lines. Two reference papers PMID 29749434 and PMID 29217166, where PMID 29217166 was verified to describe incorrect nucleotide sequence reagents (24, 26, see below), were used in PubMed similarity searches conducted in September 2019 and October 2020. Additional papers were identified through Google Scholar searches employing the keywords “gene” + “miR-145” + “cancer” conducted in April 2019 and September 2020. Publication dates were limited to 2019 to broadly align with the SGK corpus. All identified papers were visually inspected to confirm the analysis of human *miR-145* function in human cell lines.

#### Cisplatin + Gemcitabine (C+G) corpus

The cisplatin + gemcitabine (C+G) corpus included papers that described either cisplatin or gemcitabine treatment of human cancer cell lines and/or biospecimens from cisplatin or gemcitabine-treated cancer patients, where most papers also investigated human gene function. Two reference papers PMID’s 30250547 and 26852750 that both described incorrect sequences (24, 26) were used as reference papers in PubMed similarity searches conducted in September 2019 and October 2020. Additional papers were identified using Google Scholar searches with the search string “gene”, “cancer”, “cisplatin” +/− “miR” conducted in September 2019 and October 2020. A PubMed similarity search for PMID 26852750 conducted in September 2019 also identified 5 papers that referred to gemcitabine treatment (PMID’s 18636187, 26758190, 28492560, 30117016, 31272718). Four of these papers (PMID’s 26758190, 28492560, 30117016, 31272718) described incorrect sequences and were used as reference papers for PubMed similarity searches conducted in September 2019 and October 2020. Additional publications were identified through Google Scholar searches with the query “gene”, “cancer”, “gemcitabine” +/− “miR” between September 2019 and October 2020. In all cases, publication dates were limited to 2019 to align with other targeted corpora. Papers were visually inspected to confirm that they studied either cisplatin or gemcitabine treatment in the context of human cancer cell lines or biospecimens, and to exclude papers from other targeted corpora.

#### Journal corpora

*Gene* and *Oncology Reports* were selected for S&B screening as examples of journals that have published papers with incorrect nucleotide sequences (11–13), where *Oncology Reports* also published the highest number of problematic SGK papers (S1 file). Whereas *Gene* (published by Elsevier) encompasses a broad range of gene research across different species, *Oncology Reports* (published by Spandidos Publications) focusses on human cancer research.

*Gene* papers from January 2007 to December 2018 were retrieved using the Web of Science search criteria: PY= “2007-2018” AND SO= “GENE” AND DT= (“Article” OR “Review”). *Oncology Reports* papers from January 2014 to December 2018 were retrieved using the Web of Science search criteria: PY=“2014-2018” AND SO= “ONCOLOGY REPORTS” AND DT= (“Article” OR “Review”). In the case of *Gene* papers, DOI’s were retrieved, and PDF files were downloaded using the Elsevier Application Programming Interface with Crossref Content negotiation (http://tdmsupport.crossref.org), whereas open-access *Oncology Reports* articles were directly downloaded from www.spandidos-publications.com.

### Seek & Blastn screening

SGK, *miR-145* and C+G papers were named using PMID’s or journal identifiers and screened by S&B as described (12, 24, 26). All SGK papers identified for the 17 selected human genes were screened by S&B. In the case of miR-145 and C+G papers, S&B screening was conducted until 50 *miR-145*, cisplatin and gemcitabine papers were flagged for further analysis, either because S&B had flagged at least one wrongly identified reagent or had failed to extract any sequences from the text. This required S&B screening of 163 *miR-145* papers and 258 C+G papers. S&B screening was conducted in 2019 and/or 2020, with all papers flagged by S&B in 2019 being rescreened by S&B in 2020.

*Gene* papers were labelled with PMID’s, and batched pdf files were zipped into two compressed files according to publication dates (2007-2013 and 2014-2018). *Oncology Reports* papers were labelled by PMID’s and journal identifiers. S&B screening was conducted between July-October 2019, with all papers rescreened in November 2020-February 2021. *Gene* and *Oncology Reports* papers were flagged for further analysis where S&B had either flagged at least one nucleotide sequence or had failed to extract any sequences from the text.

### Visual inspection of papers following S&B screening

Papers were visually inspected to determine the claimed genetic and/or experimental identity of each sequence. If the claimed target or experimental use of any sequence was not evident, or if a sequence was claimed to target a species other than human, the sequence was excluded from further analysis. Papers that had been subject to post-publication corrections where wrongly identified nucleotide sequences had been corrected were also excluded. We included retracted papers, to align with previous descriptions of SGK papers (11–13), and in recognition of the possibility of retracted papers continuing to be cited (80).

### Manual verification of nucleotide sequence reagent identities

Nucleotide sequence identities were manually confirmed for all sequences that were not (correctly) extracted and/or flagged as being possibly incorrect by S&B, as described (26). For the *Oncology Reports* and *Gene* corpora, this involved checking at least 34% and 54% of all sequences, respectively. Further verification steps were performed for particular reagents, as follows: For reagents that were claimed to target specific gene polymorphisms or mutant sequences and for which no sequence match could be identified by either blastn or blat (25, 80), manual sequence alignments were performed in Word with the query sequence in forward, forward complement, reverse and reverse complement orientations, against either the sequence corresponding to the accession number provided within the text, or to the most relevant genomic sequence found in NCBI GenBank, according to the text claim. Sequences were delineated using the R studio “stringr” library and accepted as targeting if the specified mutated base(s), when reverted to their original base(s) as described in NCBI dbSNP https://www.ncbi.nlm.nih.gov/snp/, allowed the reagent to target the wild-type sequence according to previously published targeting criteria (12).

i. If no significant matches were identified for reagents specified for the analysis of mutant or variant targets, mismatches within the nucleotide sequence were converted to the wild type sequence, either as described in the publication or according to dbSNP and reanalysed as described (26). Reagents that were indicated to target the claimed wild-type sequence were accepted as correct targeting reagents.
ii. All flagged incorrect targeting sequences were double-checked through additional blastn searches against the database: “Homo sapiens (taxid:9606)”, optimized for “Somewhat similar sequences (blastn)”, using an expect threshold 1000, in February 2021.

Nucleotide sequence reagents that were verified to have been wrongly identified were assigned to one of 3 previously described error categories (11, 12):

i. Reagents claimed to represent targeting reagents but verified to target a human gene or target other than that claimed within the text. This error category included miR-targeting reverse RT-PCR primers with incorrect gene targeting descriptions, as supported by sequence verification (26), and by having been employed to analyse gene(s) other than the claimed miR and/or as a claimed universal primer according to Google Scholar searches (11, 13). While we recognise that some of these reagents could amplify the claimed miR target as described, their descriptions as specific targeting reagents were incorrect and could lead to incorrect RT-PCR primer reuse.
ii. Reagents claimed to target a human gene or genomic sequence but verified to be non-targeting in human. These reagents included RT-PCR primers that targeted introns or other non-transcribed regions within claimed genes.
iii. Reagents claimed to represent non-targeting reagents in human but verified to target a human gene or genomic sequence.

### Additional publication analyses

For all papers subjected to S&B analysis, publication titles were visually inspected to identify human gene identifiers, human cancer types, and drug identifiers which were confirmed through Google searches. Human genes were categorized as either protein-coding or ncRNA’s (miRs, lncRNA or circRNA) according to GeneCards (https://www.genecards.org/).

Journal publishers were identified via the SCImago database (https://www.scimagojr.com/). The Journal Impact Factor corresponding to the (closest) publication year of each paper was obtained from the Clarivate Analytics Journal Citation Reports database (https://jcr.clarivate.com/). Numbers of original articles published per year by *Gene* and *Oncology Reports* were obtained from Clarivate InCites (https://incites.clarivate.com), under Entity type= “Publication Sources”, Publication Date= 2007-2018, DT= include only “Article”. The country of origin of each paper was assigned according to the affiliations of at least half of the listed authors. Papers were considered to be affiliated with hospital(s) if the institutional affiliations of at least half of the listed authors were associated with one or more of the keywords: “clinic”, “health cent”, “hosp”, “hospital”, “infirmary”, “sanatorium”, “surgery”. Papers not meeting this criterion were considered to be affiliated with institutions other than hospitals. Proportions of problematic papers (from China versus all other countries, hospitals versus other institutions) were compared using the Fisher’s Exact test (SPSS statistics 27).

#### Bibliometric analysis of human genes in problematic papers

Linkages of protein-coding genes to publications were obtained via gene2pubmed from NCBI NIH (https://ftp.ncbi.nlm.nih.gov/gene/DATA/gene2pubmed.gz) on 15 July, 2021 as described (29, 49). Two-sided Mann Whitney U tests were performed using SciPy (82). Post-publication notices linked with problematic papers were identified through PubMed and Google Scholar searches. PubMed ID’s or other publication identifiers were employed as search queries of gene knowledgebases in May 2021 (30–34). Publication citation counts are those reported by Google Scholar in March, 2021. Problematic papers cited by clinical trials were cited by at least one publication within the NIH Open Citation collection (83), which in MedLine carried the annotation of a publication type of any “clinical trial” (without distinguishing clinical trial stage). The approximate potential to translate (APT) for problematic papers in each corpus was calculated as described (35) and obtained from iCite (83).

## Supporting information

S1 Data SGK

S2 Data miR-145

S4 Data Gene

S5 Data Oncology Reports

S5 Data Journals

## Authors’ contributions (alphabetical order)

**Conceptualization**: Jennifer Byrne, Guillaume Cabanac, Cyril Labbé, Thomas Stoeger; **Methodology:** Jennifer Byrne, Guillaume Cabanac, Cyril Labbé, Thomas Stoeger; **Formal analysis:** Jennifer Byrne, Bertrand Favier, Yasunori Park, Pranujan Pathmendra, Thomas Stoeger, Rachael West; **Writing - original draft preparation:** Jennifer Byrne, Yasunori Park, Thomas Stoeger, Rachael West; **Writing - review and editing:** Jennifer Byrne, Guillaume Cabanac, Amanda Capes-Davis, Bertrand Favier, Cyril Labbé, Yasunori Park, Pranujan Pathmendra, Thomas Stoeger, Rachael West; **Funding acquisition:** Jennifer Byrne, Amanda Capes Davis, Cyril Labbé, Thomas Stoeger; **Supervision:** Jennifer Byrne **Conflicts of Interest:** The authors have no relevant financial or non-financial interests to disclose.

## Supporting Information Captions

**S1 Fig. Problematic *Gene* papers per year, according to country of origin and institutional affiliation type.** Total numbers of problematic papers for each country/group of countries are shown in the upper left corner of each panel. A: Numbers of problematic *Gene* papers (Y axes) per publication year (X axes) according to country of origin, shown above each graph. Countries are shown in alphabetical order, from top left. B: Numbers of problematic *Gene* papers (Y axis) per publication year (X axis) from China (right panel) or all other countries (left panel). Papers affiliated with hospitals or other institution types are shown in blue or grey, respectively. Numbers of problematic papers per year are shown below each stacked bar graph.

**S2 Fig. Problematic *Oncology Reports* papers per year, according to country of origin and institutional affiliation type.** Total numbers of problematic papers for each country/group of countries are shown in the upper left corner of each panel. A: Numbers of problematic *Oncology Reports* papers (Y axes) per publication year (X axes) according to country of origin, shown above each graph. Countries are shown in alphabetical order, from top left. B: Numbers of problematic *Oncology Reports* papers (Y axis) per publication year (X axis) from China (right panel) or all other countries (left panel). Papers affiliated with hospitals or other institution types are shown in blue or grey, respectively. Numbers of problematic papers per year are shown below each stacked bar graph.

**S3 Fig. Human protein-coding genes in problematic papers appear frequently in PubMed.** Numbers (log base 10) of PubMed papers (Y axis) for primary protein-coding genes in problematic papers (A) (green) or claimed protein-coding gene targets of wrongly identified reagents (B) (green), versus all other human protein-coding genes (A, B) (grey). Horizontal lines within box plots indicate median values, with box plots showing percentiles according to letter-proportions, ie +/− 25% percentile, +/− (25 + 25/2)% percentile, +/− (25 + 25/2 + 25/4)% percentile.

**S1 Table. Wrongly identified nucleotide sequence reagents.** Wrongly identified sequences are listed for each corpus in separate tabs, as well as a combined list of all wrongly identified sequences. Primary human protein-coding and ncRNA’s refer to the first-mentioned gene in each category in each publication title.

**S2 Table. Author corrections, expressions of concern and retractions associated with problematic papers.**

**S3 Table. Problematic papers curated within gene knowledgebases.** Problematic papers identified within CancerMine, ChimerDB, emiRIT, miRTex, and mirPub are shown in separate tabs.

**S1 Data. Single gene knockdown (SGK) corpus.** All SGK papers screened by S&B and problematic SGK papers are shown in separate tabs. Primary human genes refer to the first-mentioned gene in the publication title or abstract.

**S2 Data. *miR-145* corpus.** All *miR-145* papers flagged by S&B screening and problematic *miR-145* papers are shown in separate tabs. Primary human genes refer to the first-mentioned gene(s) in the publication title or abstract.

**S3 Data. Cisplatin + gemcitabine (C+G) corpus.** All C+G papers flagged by S&B screening and problematic C+G papers are shown in separate tabs. Primary human genes refer to the first-mentioned gene(s) in the publication title or abstract.

**S4 Data. Problematic *Gene* papers.** Primary human genes refer to the first-mentioned gene(s) in the publication title or abstract.

**S5 Data. Problematic *Oncology Reports* corpus.** Primary human genes refer to the first-mentioned gene(s) in the publication title or abstract.

**S6 Data. Journals and publishers that have published problematic papers.** Journals and publishers of problematic SGK, *miR-145* and C+G papers and all problematic papers are shown in separate tabs.

