## Supplementary material for "Human gene function publications that describe wrongly identified nucleotide sequence reagents are unacceptably frequent within the genetics literature": S1 Figure

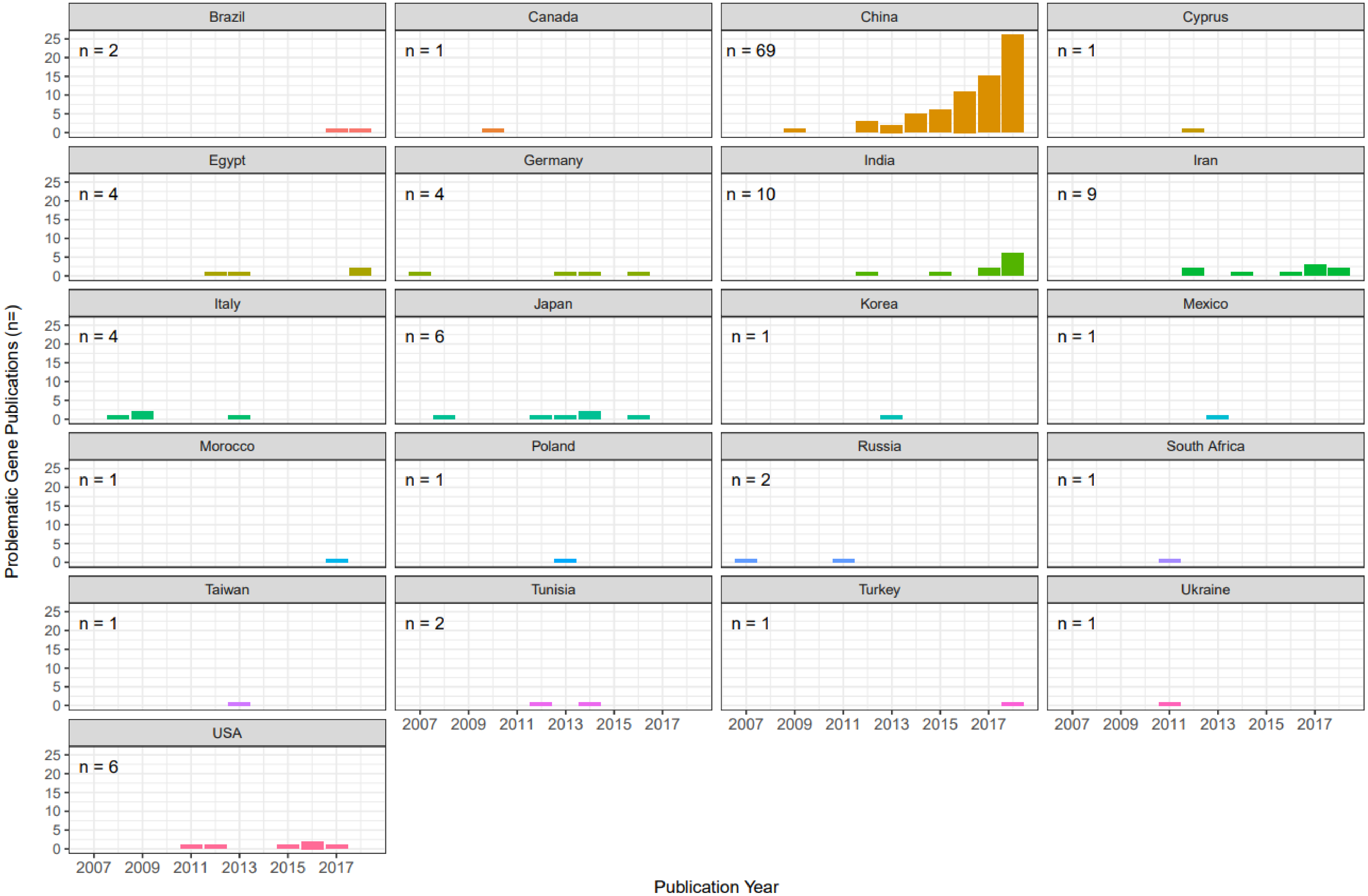


S1A Fig. Problematic Gene papers per year, according to country of origin.

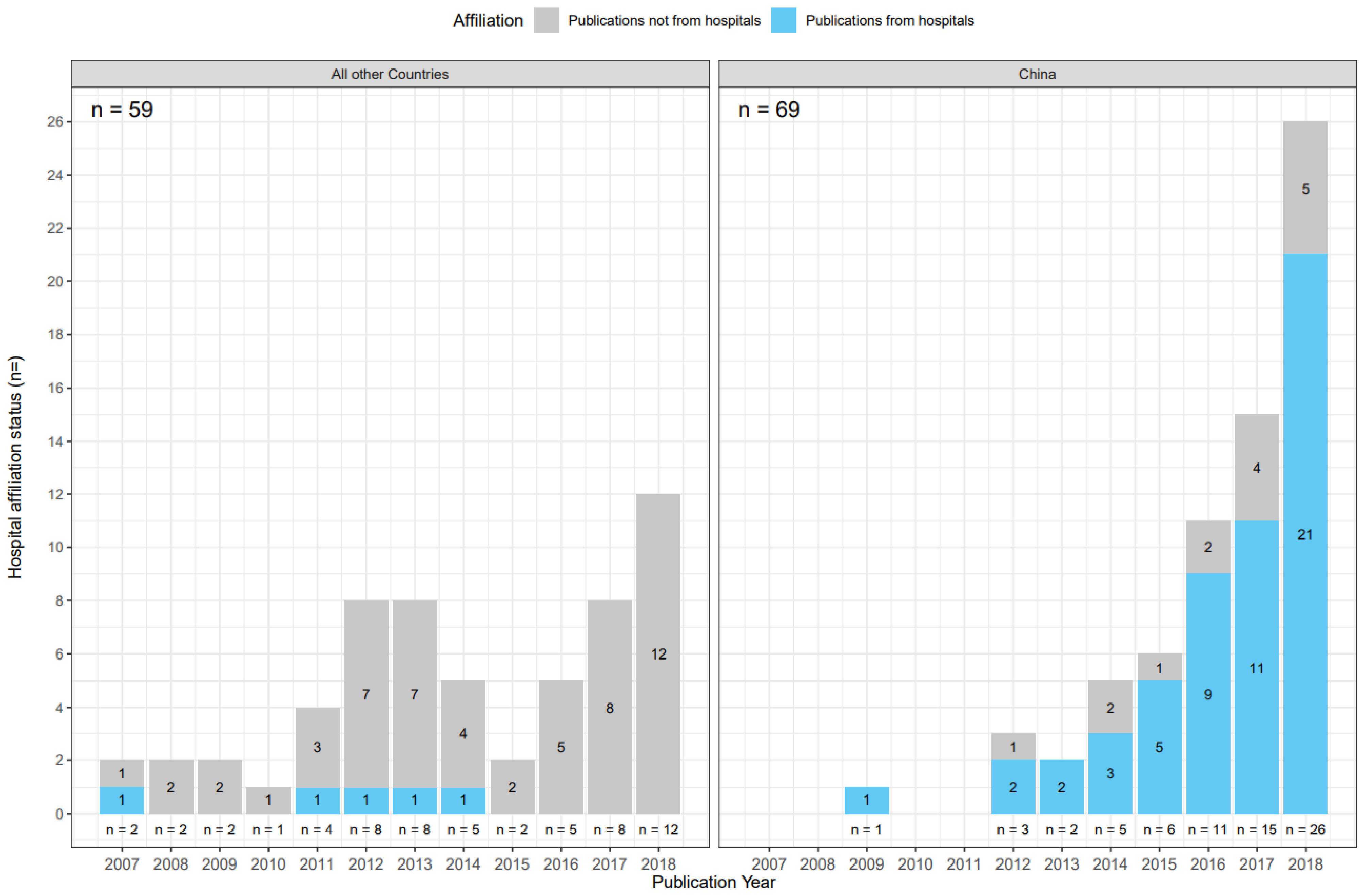


S1B Fig. Problematic Gene papers per year, according to country of origin and institutional affiliation type.
