## Supplementary material for "Human gene function publications that describe wrongly identified nucleotide sequence reagents are unacceptably frequent within the genetics literature": S2 Figure

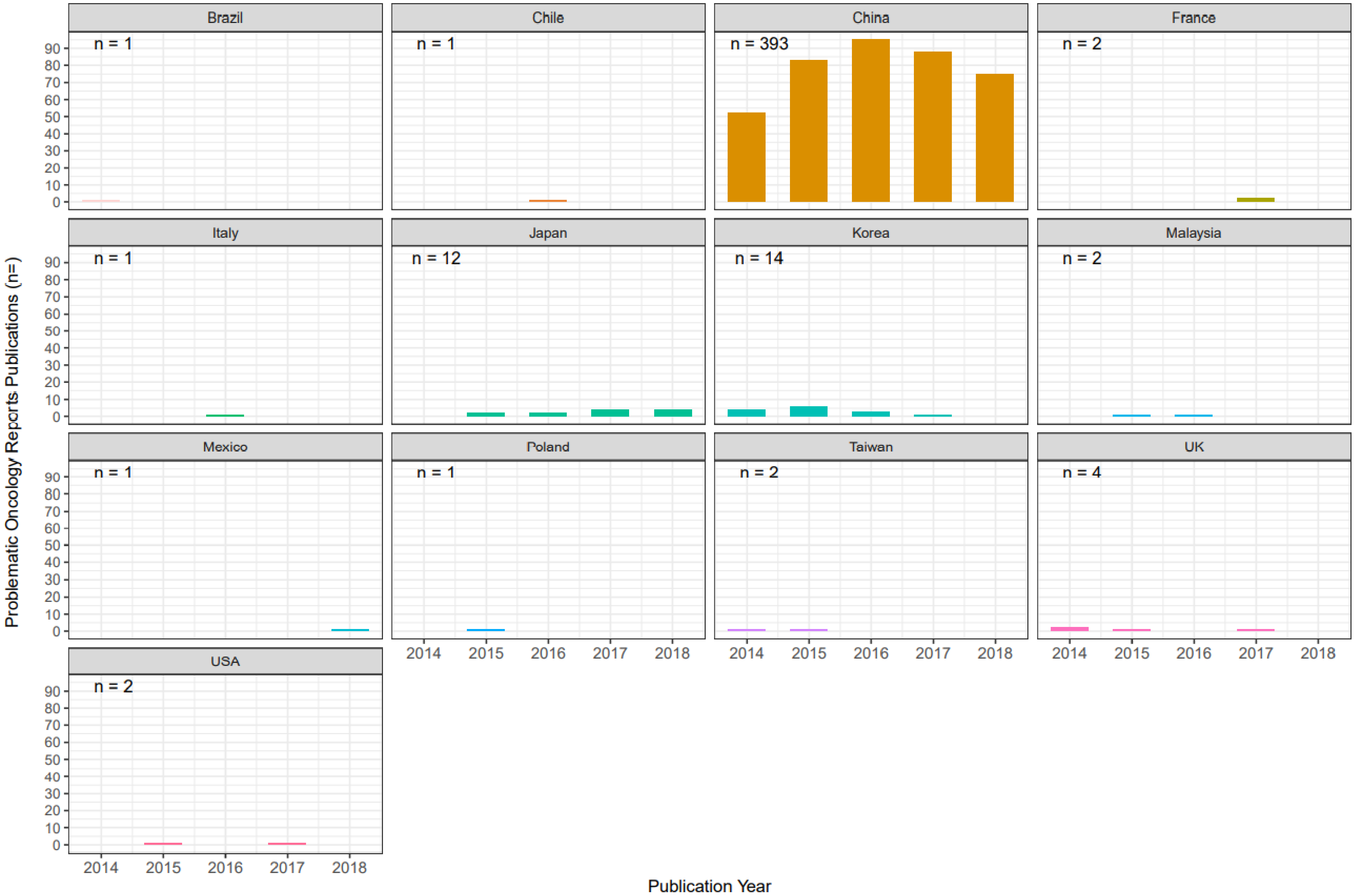


S2A Fig. Problematic Oncology Reports papers per year, according to country of origin.

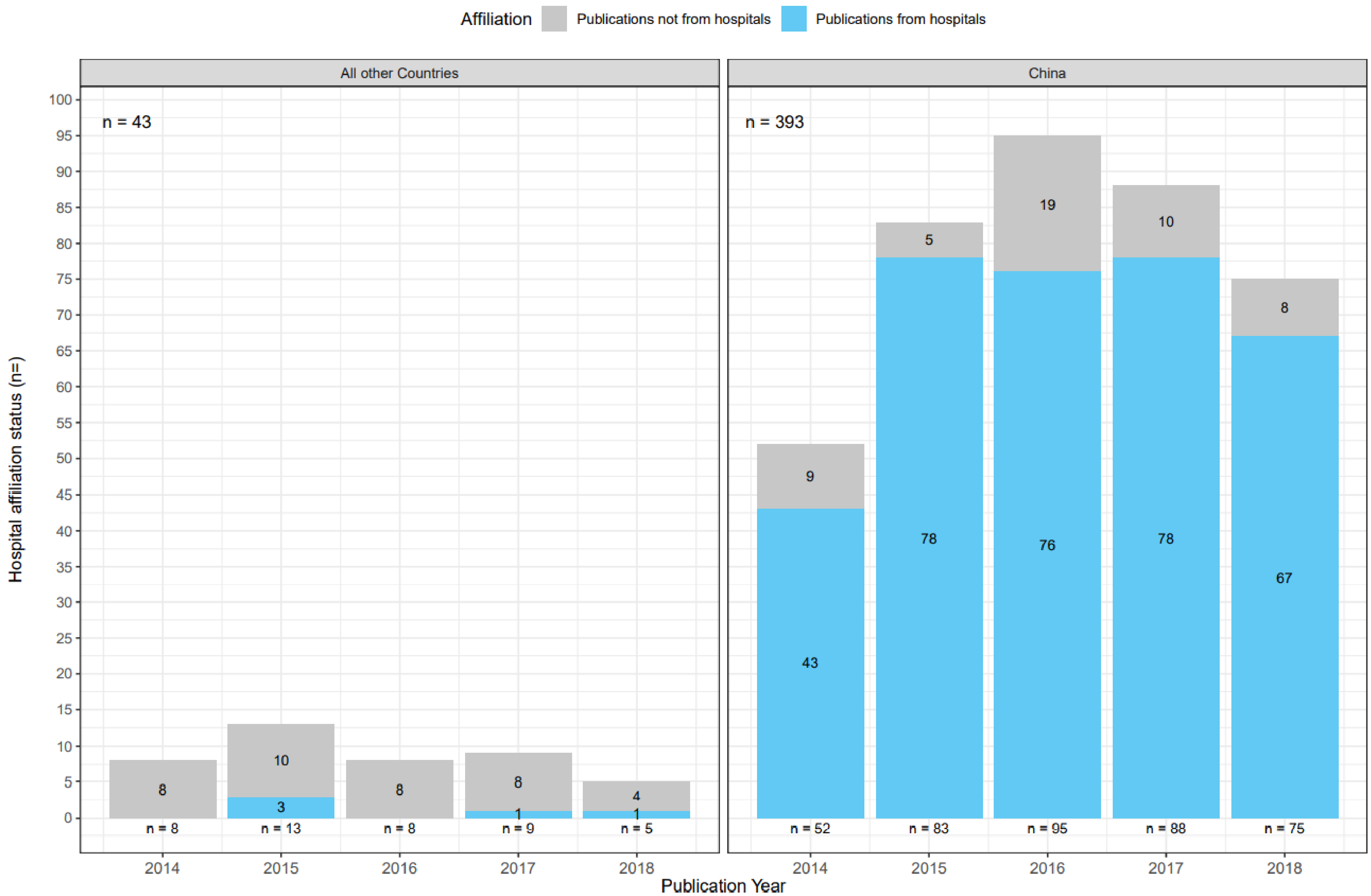


S2B Fig. Problematic Oncology Reports papers per year, according to country of origin and institutional affiliation type.
