## Supplementary figures and images for "Human gene function publications that describe wrongly identified nucleotide sequence reagents are unacceptably frequent within the genetics literature"

### S3 Figure

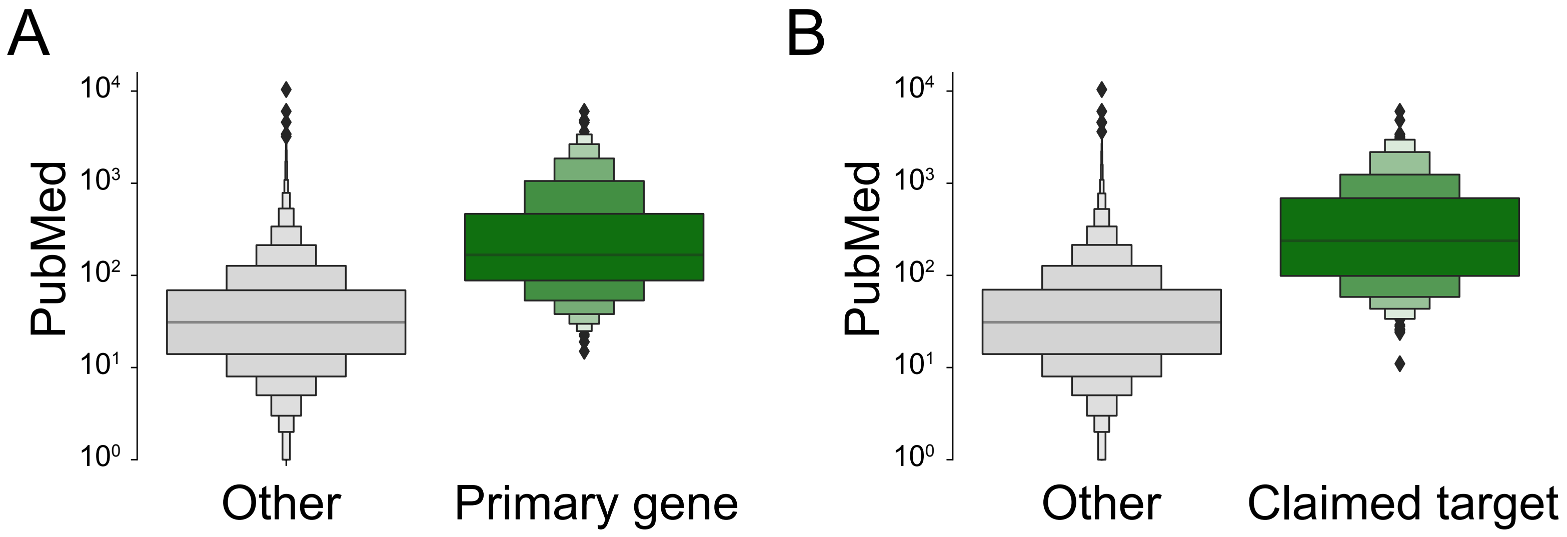


S3 Fig. Human protein-coding genes in problematic papers appear frequently in PubMed.
